# The formation of the bacterial RNA polymerase–promoter open complex involves a branched pathway

**DOI:** 10.1101/2021.03.27.437306

**Authors:** Anssi M. Malinen, Jacob Bakermans, Emil Aalto-Setälä, Martin Blessing, David L.V. Bauer, Olena Parilova, Georgiy A. Belogurov, David Dulin, Achillefs N. Kapanidis

## Abstract

The expression of most bacterial genes commences with the binding of RNA polymerase (RNAP)–σ^70^ holoenzyme to the promoter DNA. This initial RNAP–promoter closed complex undergoes a series of conformational changes, including the formation of a transcription bubble on the promoter and the loading of template DNA strand into the RNAP active site; these changes lead to the catalytically active open complex (RP_O_) state. Recent cryo-electron microscopy studies have provided detailed structural insight on the RP_O_ and putative intermediates on its formation pathway. Here, we employ single-molecule fluorescence microscopy to interrogate the conformational dynamics and reaction kinetics during real-time RP_O_ formation. We find that the RP_O_ pathway is branched, generating RP_O_ complexes with different stabilities. The RNAP cleft loops, and especially the β’ rudder, stabilise the transcription bubble. The RNAP interactions with the promoter upstream sequence (beyond −35) stimulate transcription bubble nucleation and tune the reaction path towards stable forms of the RP_O_. The mechanistic heterogeneity of the RP_O_ pathway may be a prerequisite for its regulation since such heterogeneity allows the amplification of small promoter sequence or transcription-factor-dependent changes in the free energy profile of the RP_O_ pathway to large differences in transcription efficiency.

## Introduction

Transcription initiation is the first and most regulated step in gene expression in all organisms. The expression of most bacterial genes commences with the binding of RNA polymerase (RNAP)–σ^70^ holoenzyme to the promoter DNA (1). The initial RNAP–promoter closed complex (RP_C_) undergoes large conformational changes leading to a RNAP–promoter open complex (RP_O_), which is capable of RNA synthesis. These conformational changes are of paramount importance, since their modulation by promoter DNA sequence, protein transcription factors, and small-molecule ligands strongly affects the number of active open complex, and thus the transcription efficiency (2). Further, several antimicrobials, including clinically used drugs rifampicin (3)(4) and fidaxomicin (5), exert their effect by blocking RNAP from proceeding during a specific step of transcription initiation (6). However, despite substantial progress in defining the structural basis of transcription initiation mechanism (7)(8)(9)(10), the identity, sequence, and kinetics of conformational changes leading to RP_O_ formation remain elusive.

At the initial step of the RP_O_ formation pathway, the RNAP–σ^70^ holoenzyme recognises the promoter by forming sequence-specific contacts with the −35 element, and sequence-independent contacts upstream from the −35 (“upstream sequence”) as well as around the −10 element [reviewed in (2)(11)]. In this RP_C_ state, the promoter remains fully double-stranded, but is bent by ~17° at the −10 element, thus positioning the downstream promoter DNA above the DNA-binding cleft of the RNAP (10). Studies using footprinting (12)(13)(14)(15), atomic force microscopy (16) and ensemble FRET (17) have indicated additional extensive bending and wrapping of the promoter upstream sequence (between the −35 element and −82); this bending, which is driven by the C-terminal domains of the two RNAP α-subunits (αCTDs) interacting with the promoter upstream, brings the upstream DNA to the RNAP surface, and strongly facilitates RP_O_ complex formation (18)(14)(19).

The isomerisation of the RP_C_ towards RP_O_ complex begins with the flipping of non-template DNA (ntDNA) −11 conserved adenine base from the duplex DNA to a specific pocket in σ^70^ (20) (21). The promoter melting then somehow propagates downstream until the full transcription bubble in the RP_O_ complex covers positions −11 to +2 (9). The bubble melting is coupled with the loading of downstream DNA duplex into the RNAP cleft, and the loading of single-stranded template DNA (tDNA) into the RNAP active site. Structural (9)(10) and biochemical (2) studies have identified several putative intermediates on the path from the RP_C_ to RP_O_; however, the number and structural properties of the intermediates detected appear to heavily depend on the promoter sequence, transcription factors, and experimental conditions.

The mechanism discussed above describes the formation of a uniform RP_O_ complex on a standard linear reaction pathway. A more complete description of the transcription initiation, however, needs to consider several studies that suggested that individual RP_O_ molecules are not identical, and they instead differ in functional properties (22)(23)(24)(25). One of the most notable variation among RP_O_ complexes is their tendency to perform abortive initiation, i.e., the premature release of short RNAs synthesised by promoter-bound RNAP [reviewed in (26)]. In fact, it has been estimated that >50% of the RP_O_ complexes are permanently locked into the abortive initiation mode and cannot produce full-length RNA (22)(23)(24)(25). The presence of at least two different RP_O_ classes – one productive and one non-productive (abortive) – raises the possibility that the RP_O_ pathway is also not linear, but instead branches to allow the formation of structurally and functionally different RP_O_ molecules. It has been further suggested that the ratio of productive and non-productive RP_O_ complexes can be modulated by transcription factors and thus offers a layer for gene regulation in the cell (27). On the other hand, recent single-molecule studies revealed long-lived pausing, backtracking and arrest of initially transcribing bacterial and mitochondrial RNAPs that could potentially explain the productive and abortive RNA synthesis by a single type of RP_O_ complexes (28)(29)(30). The RP_O_ formation pathway branching – its occurrence and mechanism – thus warrants further study.

Here, we utilise single-molecule techniques to complement bulk biochemical assays and resolve asynchronous, multi-step and branched reaction mechanisms during σ^70^-dependent RP_O_ formation. Our results demonstrate that the RP_O_ formation pathway is indeed branched, resulting in RP_O_ complexes with markedly different stability in their open transcription bubble conformation. Furthermore, αCTD interactions with the promoter upstream sequence strongly stimulate bubble initiation and tune the reaction pathway towards more stable RP_O_ complexes. The RNAP cleft loops (and especially the β’ rudder one), play a key role in stabilising the open transcription bubble. We propose that the inherent mechanistic heterogeneity of the RP_O_ pathway is a prerequisite for transcriptional regulators to strongly modulate the formation and function of RP_O_ complexes, and the rate of transcript generation.

## Methods

### Promoter preparation

Labelled and unlabelled DNA oligos to make *lacCONS+2* promoter constructs were purchased from IBA Lifesciences (Germany) (**Fig. S1A-C**). Short *lacCONS+2* promoters (−39/+25) were reconstituted by annealing PAGE purified labelled template and non-template DNA oligos at 1 μM in annealing buffer [10 mM Tris-HCl (pH 8.0), 50 mM NaCl, 0.1 mM EDTA]. The annealing program consisted of initial denaturation (93°C, 3 min) followed by step-wise cooling to 4°C (each step: 1.2°C, 30 s). DNA strands for the long *lacCONS+2* promoters (−89/+25) were constructed using a previously described protocol (29).

### Protein preparation

*Escherichia coli* core RNAPs were expressed in *E. coli* and purified as previously described (31). The wild-type RNAP was expressed from plasmid pVS10 (T7p-α-β-β’_His_6_-T7p-ω) (32). ΔRudder loop RNAP (T7p-α-β-β’[ΔN309-K325]_TEV_His_10_-T7p-ω), Δlid loop RNAP (T7p-α-β-β’[ΔP251-S263→GG]_TEV_His_10_-T7p-ω) and Δgate loop RNAP (T7p-α-His_6__β[ΔR368-P376→GG]-β’-ω) were expressed from pMT041, pHM001 and pTG011, respectively (33). Wild-type *E. coli* σ^70^ was purified as previously described (34). Holoenzymes were assembled by incubating 0.5 μM RNAP with 1.5 μM σ^70^ for 15 min at 30°C in RNAP storage buffer [20 mM Tris-HCl (pH 8.0), 150 mM NaCl, 50% (vol/vol) glycerol, 0.1 mM EDTA, 0.1 mM dithiothreitol (DTT)].

### Microscope coverslip preparation

Borosilicate glass coverslips (1.5 MenzelGläzer, Germany) were heated to 500°C in oven for 1 h to reduce background fluorescence. The coverslips were then rinsed with HPLC-grade acetone and immerged into 1% (v/v) Vectabond (product code #SP-1800, Vector Labs, CA, USA) in acetone for 10 min to functionalise the glass surface with amino groups. Coverslips were then rinsed with acetone followed by deionized water before drying them under a stream of nitrogen gas. A silicone gasket (103280, Grace Bio-Labs, OR, USA) with four reaction wells was placed in the middle of the coverslip. The coverslip surface was then simultaneously passivated by pegylation against unspecific protein/DNA binding and biotinylated to provide attachment points for specific protein immobilisation. Each well on the coverslip was thus filled with 20 μl of 180 mg/ml methoxy-PEG (5 kDa)-SVA (Laysan Bio, AL, USA) and 4.4 mg/ml biotin-PEG (5 kDa)-SC (Laysan Bio, AL, USA) in 50 mM MOPS-KOH buffer (pH 7.5), incubated for ~3 h at room temperature and finally the wells were thoroughly rinsed with phosphate-buffered saline (PBS; Sigma Aldrich, UK). The coverslips remained functional for at least two weeks when stored at 4°C in plastic pipette tip box containing a layer of deionised water at the bottom. During the storage the coverslip wells were filled with PBS.

### Single-molecule experiments

On the day of microscopy experiment, the pegylated coverslips were incubated for ~ 10 min with 0.5 mg/ml of Neutravidin (31050, ThermoFisher Scientific, UK) in 0.5 × PBS and subsequently rinsed with 1 × PBS. The coverslips were then incubated for ~ 10 min with 4 μg/ml of Penta•His biotin conjugate antibody (34440, Qiagen, UK) in reaction buffer [40 mM HEPES buffer (pH 7.3, BP299100, Fisher Scientific, UK), 100 mM potassium glutamate, 10 mM MgCl_2_, 1 mM DTT, 1 mM cysteamine hydrochloride, 5% glycerol (vol/vol), 0.2 mg/ml bovine serum albumin] and subsequently rinsed with the reaction buffer.

To analyse the RP_O_ complex formation in real-time at 22°C the anti-His-tag-antibody coated coverslip was incubated ~10 min with 1 nM label-free holoenzyme in the reaction buffer, rinsed thoroughly with the reaction buffer and mounted on the microscope. 25 μl of imaging buffer [i.e., reaction buffer supplemented with 2 mM UV-treated Trolox, 1% (w/v) glucose, 0.4 μg/ml catalase (10106810001, Roche Diagnostics, Germany), 1 μg/ml glucose oxidase (G2133, Sigma Aldrich, UK)] was replaced to the imaged well. Data recorder was started to take an 80 s movie. 1 μl of 4 nM promoter in the reaction buffer was gently pipetted to the well at ~ 8 s time-point. The formation of RNAP–promoter complexes was evident by the appearance of bright co-localised spots on the Cy3B and ATTO647N fluorescence channels. In some experiments these surface-formed RNAP–promoter complexes were further imaged after exchanging fresh imaging buffer to the well and finding non-bleached field-of-view. The age of RNAP–promoter complexes at the moment of recording these 20 s post-binding movies was ~ 3–7 min. In some control experiments, we monitored the initial RNA synthesis activity RNAP by including 1 mM NTPs (ATP, GTP, CTP and UTP) in the imaging buffer.

To analyse transcription bubble dynamics in pre-assembled RP_O_ complexes, 2 nM holoenzyme was incubated with 5 nM promoter in reaction buffer for 15 min at 37°C. 100 μg/ml sodium heparin (H4784, Sigma, UK) was added to disrupt non-specific RNAP–promoter complexes and ~ 1.3 μl of the mixture was transferred to anti-His-tag-antibody coated coverslip well containing 25 μl reaction buffer. The RP_O_ complex immobilisation at 22°C was let to continue until ~50 molecules were detected on the field-of-view. The well was then rinsed with reaction buffer and finally filled with 25 μl imaging buffer. Data was recorded as 20 s movies from about ten field-of-view per well at 22°C.

Single RNAP–promoter complexes were imaged using objective-based single-molecule TIRF microscope previously described (35). The donor (Cy3B) and acceptor (ATTO647N) fluorophores in the promoter were excited using 532 nm and the 642 nm lasers in alternating laser excitation (ALEX) mode, respectively (36). The emission of donor and acceptor fluorophores was separated from each other and from the excitation light, using dichroic mirrors and optical filters, and recorded side-by-side on an electron multiplying charge-coupled device camera (iXon 897, Andor Technologies, Northern Ireland). The frame time of the recordings was 20 ms with 10 ms ALEX excitation by each laser. The measured laser power before the dichroic mirror was set to ~ 4 mW and ~ 1 mW for the 532 nm and 642 nm laser, respectively.

### Single-molecule data analysis

The recorded movies were processed with custom-made TwoTone TIRF-FRET analysis software (35) (https://groups.physics.ox.ac.uk/genemachines/group/Main.Software.html) to identify and extract the intensities of co-localised donor and acceptor fluorophores. If the processed movies had fluorescent complexes on the surface already at the beginning of the movie, i.e., post-binding and pre-formed RP_O_ complex movies (see above), the following Twotone-ALEX parameters were applied to select only complexes containing a single ATTO647N acceptor dye and a single Cy3B donor dye: channel filter as DexDem&&AexAem (colocalisation of the donor dye signal upon donor laser excitation, the acceptor dye signal upon acceptor laser excitation), a width limit between the donor and the acceptor between 1 and 2 pixel, a nearest-neighbour limit of 6 pixels, and signal averaging from the frames 2–40. In the case of real-time RP_O_ complex formation movies, the nearest-neighbour limit was turned off and the time-window for the search of the surface-bound fluorescent molecules was set with the signal averaging setting (i.e. typically frames ~ 1000–3000) to the part of the movie in which most promoter binding events took place. The trajectories selected by the TwoTone-ALEX analysis were manually sorted by eliminating all traces that displayed extensive fluorophore blinking, multi-step photobleaching indicating more than one donor or acceptor dye in the same diffraction limited intensity spot, or other aberrant behaviour.

The apparent FRET efficiency (E*) was calculated using **Equation 1** where *I*_DD_ and *I*_DA_ are the emission intensities of the donor (Cy3B) and acceptor (ATTO647N) dyes upon donor excitation (532 nm), respectively (37).

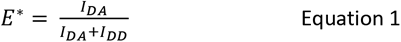

The trajectories were analysed using a modified version of the hidden Markov model ebFRET software (38) (29) (the modified code is available from the corresponding authors on reasonable request). The trajectories from the pre-formed RP_O_ or post-binding movies were fit to two-state HMM model followed by noise filtering by requiring an accepted dwell time to satisfy the criteria that the step (i.e., change in E*) is separated from the subsequent step by more than 3-fold the Allan deviation (39) (29). The trajectories were then classified into dynamic or static populations depending whether they displayed >2 or ≤2 accepted E* transitions, respectively. The dwell times were extracted from the dynamic trajectories to compile a dwell time distribution. The dwells with undefined length, i.e. the first and last dwell, were discarded at this point.

The trajectories from the real-time RP_O_ formation movies were analysed separately for the first transcription bubble opening event, i.e., the RP_C_→RP_O_ transition, and transcription bubble dynamics after the RP_O_ formation. The latter analysis was identical to the case of pre-formed RP_O_ complexes with the exception that the RP_C_→RP_O_ transition at the beginning of the trajectory was trimmed away before HMM. In contrast, the analysis of the RP_C_→RP_O_ transition in the trajectories was performed after trimming away possible bubble dynamics subsequent to the first transcription bubble opening event. We fit the first bubble opening trajectories using two-state HMM, i.e., RP_C_→RP_O_ mechanism, and three-state HMM, i.e., RP_C_→RP_i_→RP_O_ mechanism. The initial fits were filtered by requiring true state transitions to be at least 2-fold the Allan deviation (39) (29). Selection of the more complex 3-state model for the trajectory also required that both the HMM lower bound value and Aikake information criteria, calculated as previously described (40), favoured this model. The dwell times in the RP_C_ and RP_i_ states were compiled to separate dwell time distributions. The lifetime of the RP_C_ state was determined by fitting the dwell time distribution to the mono-exponential decay function using Origin software (OriginLab Corporation, MA, USA). Due to the limited amount of data, the lifetime of RP_i_ state was calculated as the arithmetic mean and its 95% confidence interval was estimated by bootstrapping (10000).

The histograms of *E*\* values were fit in Origin software to one or two Gaussian distributions using **Equation 2** with *n* fixed as 1 or 2, respectively. The fit parameters *E*_*c*_ ^*^, *w* and *A* are the centre, width and area of the Gaussian distributions, respectively.

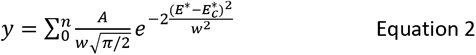

## Results

### Direct formation of surface-immobilised catalytically active open complexes

To be able to monitor RNAP–promoter open complex (RP_O_) formation in real-time at the single-molecule level, we used FRET to look at the changes in distances between two points, i.e., positions −15 and +15 relative to the transcription start site (position +1) on a promoter DNA fragment. A fluorophore pair incorporated in positions −15 (donor) and +15 (acceptor) produces FRET signatures that vary depending on the transcription bubble conformation; this pair has been employed before to monitor conformational changes in populations of single transcription complexes (41)(42), conformational dynamics of RP_O_ complexes (43) and conformational changes after the formation of RP_O_ complex (29) on a consensus *lac* promoter (*lacCONS+2*) (**Fig. S1A,B**).

Here, we modified our previous protocols to detect the nascent RNAP–promoter complex (RP_C_) and its subsequent maturation to RP_O_ (**Fig. 1**). To this end, we attached molecules of the *Escherichia coli* RNA polymerase–σ^70^ holoenzyme to the surface of a coverslip and started imaging the surface using TIRF microscopy (**Fig. 1A,B**). Subsequent addition of the dual-labelled promoter DNA to the reaction buffer was expected to lead to the appearance of co-localised fluorescent spots on the donor (Cy3B label) and acceptor (ATTO647N label) detection channels of the microscope upon binding to the surface-attached holoenzyme (**Fig. 1B**). The timing of the appearance of the fluorescent spots on the surface (due to DNA binding and formation of RP_C_ complexes) is precisely defined in the single-molecule trajectories by the simultaneous appearance of Cy3B and ATTO647N fluorescence signals (“DNA binds” time point, **Fig. 1C**). The −15/+15 ruler reports low FRET for the RP_C_ complex, and intermediate FRET for the RP_O_ complex, since the formation of the transcription bubble decreases the distance between the −15 and +15 positions in the DNA (43). The RP_C_ → RP_O_ transition in the trajectories is thus indicated by a sharp FRET increase (“DNA transcription bubble opens” time point, **Fig. 1C**), which may occur in one or several steps, depending on the intermediates on the reaction pathway.

**Figure 1.**
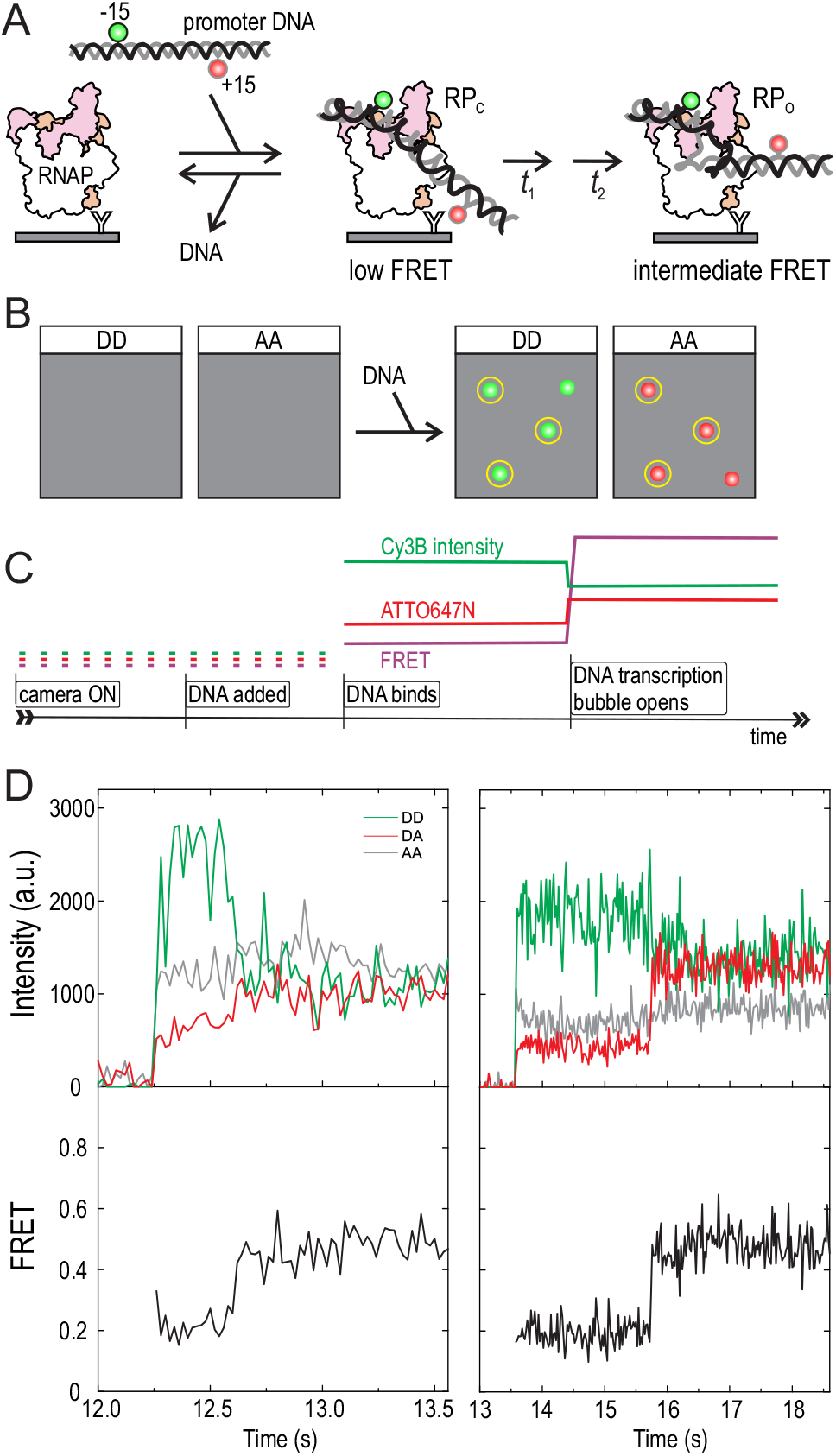
Single-molecule FRET method to monitor the RNAP–promoter open complex formation in real-time. (A) *E.* coli RNAP–σ^70^holoenzyme is immobilised on the PEGylated microscope coverslip using biotinylated anti-His-tag-antibody. *lacCONS+2* promoter, which is labelled with a donor fluorophore (D, Cy3B) at non-template DNA position −15 and an acceptor fluorophore (A, ATTO647N) at template DNA position +15, is added to the reaction buffer. The promoter binds to the RNAP and becomes visible on the coverslip surface. The initial RNAP–promoter closed complex isomerises to the open complex, which decreases the distance between the −15 and +15 dyes and increases the FRET. (B) Schematic microscope field-of-view before and after promoter addition to the reaction buffer. Data on the DD (donor excitation–donor emission) and AA (acceptor excitation–acceptor emission) channels is used to identify the RNAP–promoter complexes containing both the Cy3B and ATTO647N dyes. These molecules are highlighted with yellow circles. (C) Schematic single-molecule trajectory showing promoter binding to the RNAP and subsequent isomerisation to the open complex state. Abrupt increase in the Cy3B and ATTO647N fluorescence intensities defines the moment of promoter binding. The increase in the FRET from the low level to the intermediate level defines the moment of open complex formation. (D) Two experimental trajectories show promoter binding and open complex formation. DA indicates the signal on the donor excitation–acceptor emission channel.

Our experimental single-molecule trajectories indeed show the expected fluorescence intensity and FRET signatures of RP_C_ complex formation and isomerisation to the RP_O_ state (**Fig. 1D**). The moment of RNAP-promoter complex formation was precisely defined by the simultaneous appearance of Cy3B and ATTO647N fluorescence in the single-molecule trajectories (e.g., at 12.25s and 13.5s in the left and right panels, respectively, of **Fig. 1D**). The apparent FRET efficiency (E*) of the first stable complexes was E*~0.2 (**Fig. 1D**), a value identical to that we obtained previously for the closed transcription bubble state (43). After a short time (~0.35s and ~2s in the traces of **Fig. 1D**), the FRET increased to E*~0.45, a value identical to that we obtained previously for the open transcription bubble state (43). DNA binding to the coverslip surface was strictly mediated by the RNAP, since the number of non-specific DNA binding events was negligible in the absence of RNAP on the surface (cf. **Fig. S2A** to **S2B**). On the population level, the newly formed RNAP–promoter complexes displayed a bimodal FRET distribution, with mean FRET values ∼0.2 and ∼0.45 (**Fig. S2C**) contrasting with the unimodal FRET distribution (mean ~0.18) of the protein-free immobilised promoter DNA (**Fig. S2D**). To test whether the ∼0.45 FRET state is indeed a catalytically competent RP_O_ complex, we added NTPs to the sample buffer; this addition almost eliminated the ∼0.45 FRET state, as expected if RP_O_ complexes engage RNA synthesis and escape the promoter (**Fig. S2C**).

To provide further proof for the formation of catalytically active RP_O_ complexes *in situ* on the coverslip surface, we performed experiments using a promoter with a different labelling scheme, which is very effective in monitoring the progress of initial transcription (dyes at positions −15 and +20; **Fig. S1C**). The scrunching of the downstream DNA towards the RNAP during initial RNA synthesis leads to a stepwise increase in FRET until RNAP escape from the promoter returns the FRET to a low level (**Fig. S3A,B)** (24). Example trajectories in **Fig. S3C** demonstrate abortive initiation and promoter-escape events occurring shortly after the formation of the RNAP–promoter complexes. However, we note that some RNAP–promoter complexes (typically 20–50% of all complexes) on the surface neither form RP_O_ nor engage RNA synthesis; these molecules remain in stable low FRET state (~0.2) and may thus represent unproductive complexes resulting, e.g., from RNAP binding to the ends of the promoter DNA fragment (**Fig. S3D**). Because the FRET sensitivity of the −15/+15 or −15/+20 labelled promoters is not sufficient to confidently distinguish RP_C_ complex from non-specific RNAP–DNA complexes, we decided to analyse further only the RNAP–promoter complexes which directly show the appearance of the FRET signature of the RP_O_ complex (i.e., the ~0.45 FRET state) on the −15/+15 labelled promoter.

### Extended promoter upstream sequence stimulates RP_O_ formation

To study the kinetics of RP_O_ complex formation in real-time and its modulation by the αCTD–promoter upstream interactions, we performed experiments using a long (DNA extending from position −89 to position +25) and short (DNA extending from position −39 to position +25) version of the *lacCONS+2* promoter (**Fig. 2A**). We also examined the kinetics of open complex formation by using additional versions of the promoter DNAs, which were either fully double-stranded (dsLC2 promoter; **Fig. S1A**) or contained a mismatch in the promoter region from −10 to −4 (a.k.a. pre-melted promoter, pmLC2; **Fig. S1B**).

**Figure 2.**
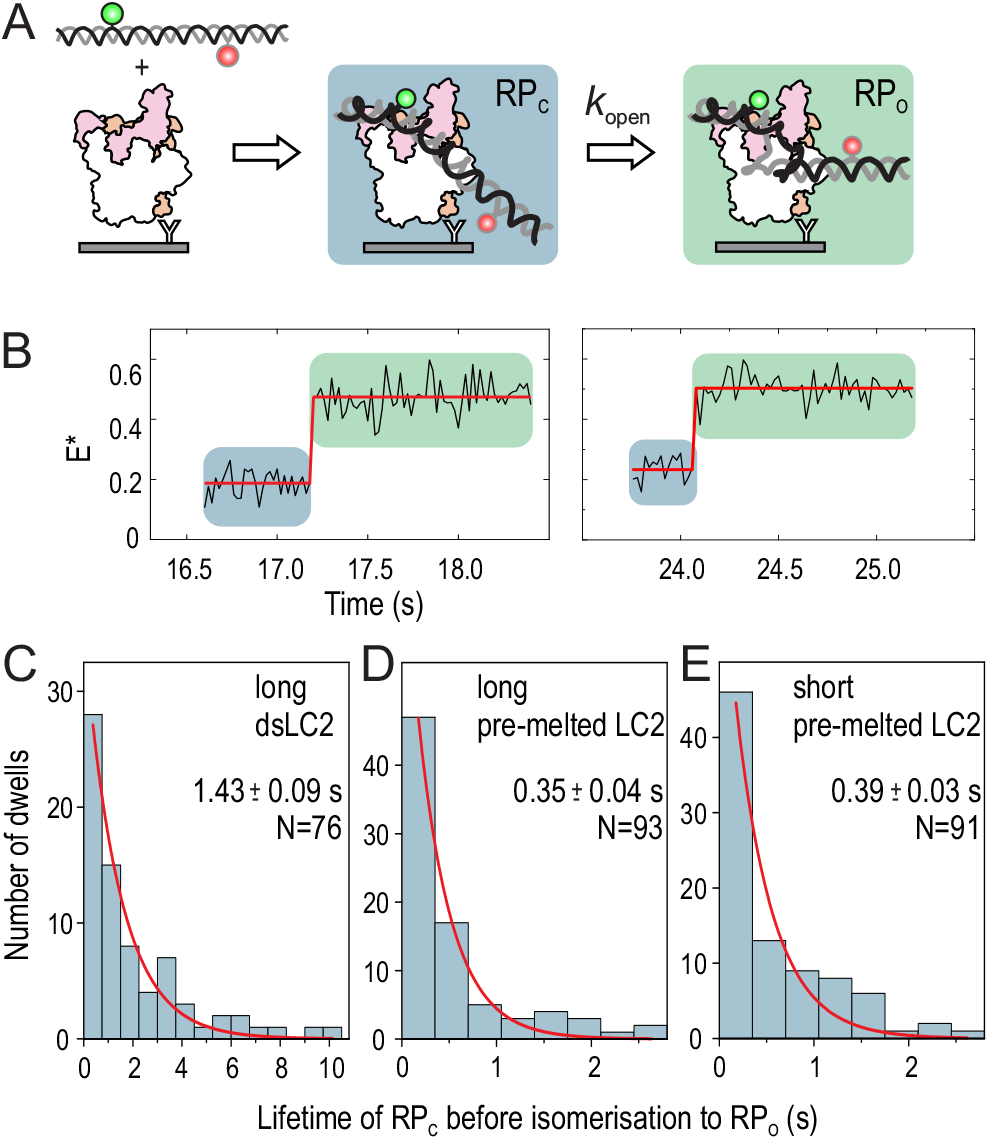
Rate of RP_O_ formation. (A) Schematic representation of the real-time RP_O_ formation experiment. The promoter has donor and acceptor labels at positions −15 and +15, respectively. (B) Example trajectory on the left demonstrates promoter binding to the surface-immobilised RNAP and the formation of RP_C_ complex at 16.6 s. The RP_C_ isomerises to RP_O_ complex at 17.2 s. Another example trajectory of real-time RP_O_ complex formation is shown on the right. Dwell-times in the RP_C_ state were fit to mono-exponential function to obtain the rate constant of RP_O_ complex formation (C) on the long dsLC2 promoter, (D) long pre-melted LC2 promoter and (E) short pre-melted LC2 promoter.

RP_O_ formation was inefficient in the case of short dsLC2 at 22 °C; in fact, we could identify only 5 real-time promoter-binding events indicating RP_O_ complex formation (3% of all promoter-binding events; **Fig. 2B**); even after prolonged incubation of the RNAP–promoter complexes (~5 min) on the surface, the RP_O_ complex (i.e., the FRET species with E*~0.45) was nearly absent from the population histogram (**Fig. S4A**). In contrast, the RP_O_ complex formed efficiently at 22°C on the long promoters, as well as on the short pre-melted promoter, as seen in the E* histograms (**Fig. S4B,C,D**) and individual trajectories (**Fig. 2B**).

We then performed Hidden Markov modelling (HMM) of the trajectories to extract the dwell times in the RP_C_ state (E*~0.2) before transcription bubble opening and RP_O_ complex formation (**Fig. 2B**). The observed distribution of dwell times in the RP_C_ state for the long dsLC2 promoter (**Fig. 2C**) was fitted to a mono-exponential decay function to determine a mean lifetime for the RP_C_ complex of 1.43±0.09 s (±SE). We also tried to fit the dwell-time distribution using a double-exponential equation, but rejected this more complex kinetic model because the fit parameters were poorly defined as evident from large SE. Using a similar analysis, we estimated the RP_C_ complex lifetime as 0.35±0.04 s on the long pre-melted LC2 (**Fig. 2D**) and 0.39±0.03 s on the short pre-melted LC2 (**Fig. 2E**), respectively. These values indicate that the αCTD interactions with the upstream sequence (−89 to −40) significantly enhance the isomerisation rate of the RP_C_ to RP_O_ complex; however, this happens only on a fully double-stranded promoter. Because the introduction of the pre-melted region (−10/−4) to the promoter nearly equalised the rate of RP_C_ isomerisation to the RP_O_ complex on the short and long promoters, the αCTD–promoter interactions appear to predominantly contribute to the lowering of the activation energy of initial transcription bubble nucleation.

### A subpopulation of RP_O_ complexes form via a kinetically significant intermediate

Close inspection of the HMM fit to the RP_O_ formation FRET trajectories revealed that, even though most bubble-opening events were described by a two-state model, (i.e., the promoter conformation in the initial complex changed to the RP_O_ state in a single step; **Fig. 2B**), a subpopulation included an intermediate state (hereafter, RP_i_ complex) between the initial RP_C_ and final RP_O_ states (**Fig. 3A**).

**Figure 3.**
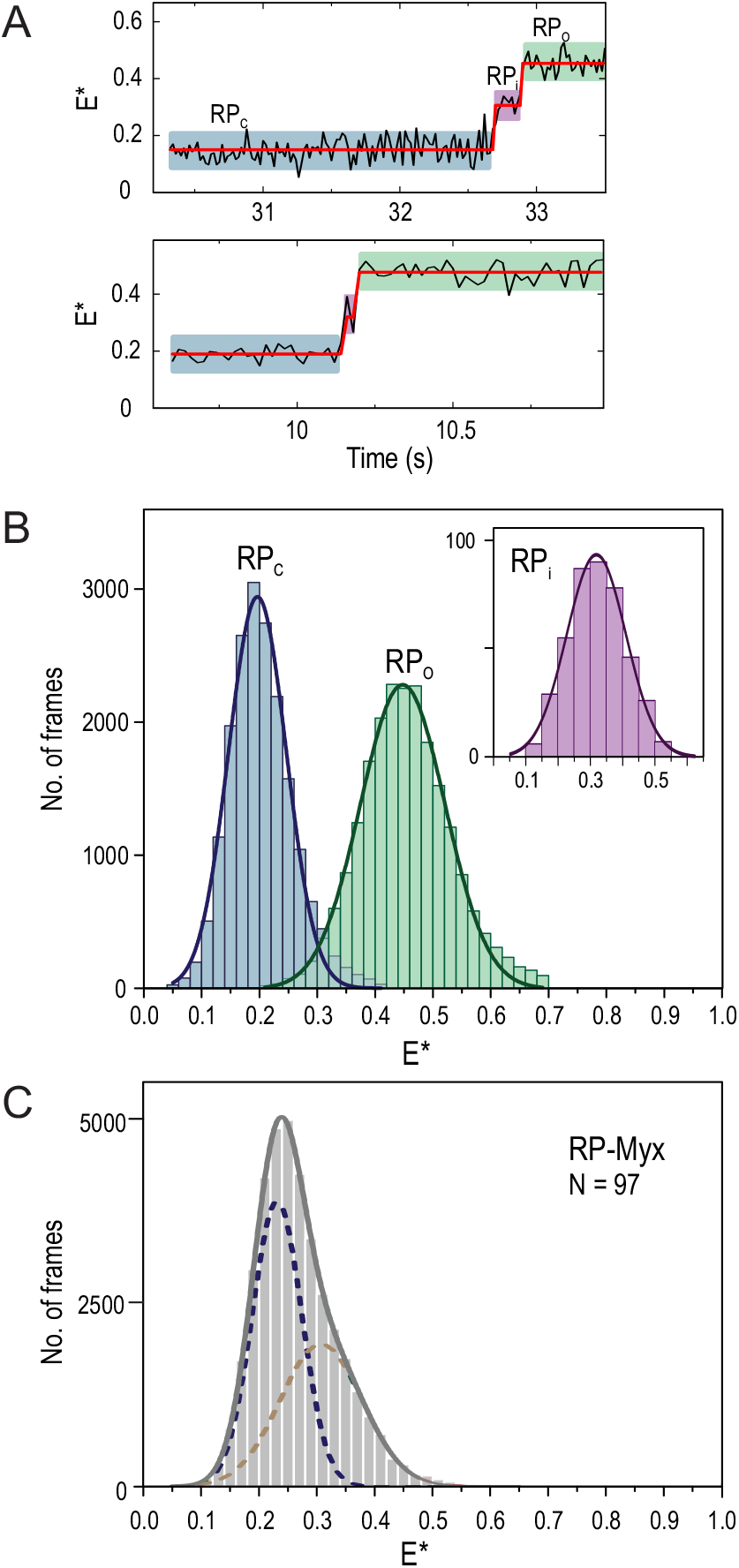
Intermediate on the RP_O_ complex formation pathway. (A) Example trajectories demonstrate the presence of an intermediate, RP_i_ complex, on the pathway from RP_C_ to RP_O_ complex. The trajectory was fit to three state HMM. (B) E* histograms for the RP_C_, RP_i_ and RP_O_ states were extracted from the HMM segmented trajectories. The E* distributions were fit Equation 2. Data from different promoter versions was pooled. (C) E* histogram of the RNAP–promoter complexes formed in the presence of 100 μM Myx inhibitor. The complexes were imaged ~5 min after their initial formation on the coverslip surface. E* distribution was fitto Equation 2.

Specifically, the RP_i_ complex was identified in 20% (exact 95% binomial confidence interval (44): 11– 30%), in 14% (8–23%) and in 14% (8–23%) of all trajectories in the case of long dsLC2, long pmLC2 and short pmLC2 promoter, respectively. The arithmetic mean lifetime of the identified RP_i_ state was 0.32 s (95% CI: 0.18–0.49 s, N=15), 0.15 s (0.07–0.26 s, N=13) and 0.17 s (0.10–0.23 s, N=13) on the long dsLC2, long pre-melted LC2 and short pre-melted LC2 promoter, respectively. Estimated lifetimes overlap within the confidence interval, suggesting that the lifetime does not depend strongly on the used promoter type. Further, the lifetime estimates indicate that the absence of the RP_i_ state in most trajectories is not explained by our existing temporal resolution (20 ms per frame); even in the worst-case scenario (assuming the shortest lifetime, 0.07 s, within the CI and assumed single exponential dwell distribution), the RP_i_ state would be missed only in 25% of the trajectories. The expected missing rate drops to only 6% when the mean RP_i_ lifetime on the long dsLC2 promoter (0.32 s) is used in the calculations.

To compare the mean FRET of the RP_i_ intermediate to that of the RP_C_ and RP_O_ complexes, we extracted FRET efficiency histograms from HMM-segmented trajectories for each of the three states (**Fig. 3B**). The mean FRET values, obtained as the centres of the fit Gaussian distribution (Equation 2), were found as 0.196±0.001 (±SE), 0.318±0.002 and 0.448±0.001 for the RP_C_, RP_i_ and RP_O_ complex, respectively. The mean FRET values suggest that the average distance between the −15 and +15 labels in the RP_i_ state has become shorter than in the RP_C_ complex but remains still longer than in the mature RP_O_ complex.

To probe the structural origin of the RP_i_ state, we included in the reaction buffer the RNAP inhibitor myxopyronin B (Myx). Biochemical and structural studies using Myx (45) and structural studies using corallopyronin A (Cor) (9), a Myx analogue, have suggested that this class of RNAP inhibitors block the formation of RP_O_ complex by preventing the loading of template DNA strand into the active site cleft. We observed that the FRET distribution of the RNAP–promoter complexes preformed in the presence of Myx was described by two Gaussians with mean FRET values 0.231±0.001 and 0.307±0.010 (**Fig. 3C**, long dsLC2 promoter). The inspection of individual trajectories revealed three classes of molecules: the first and most abundant class involved RNAP–promoter–Myx complexes characterised by an E* ~0.3 state (N = 58, 60% of all molecules, **Fig. S5A**). Interestingly, a sub-fraction (N = 19/58) of these molecules stochastically sampled a very short-lived, i.e., typically 20–40 ms (1–2 frames), higher E* state (**Fig. S5B**). The second class of molecules involved potential non-productive complexes as indicated by a stable E* ~0.2 value (N = 37, 38%, **Fig. S5C**) and the third class involved RP_O_ complexes characterized by a long-lived E* ~0.45 state (N = 2, 2%, **Fig. S5D**). Preformed complexes on the long pre-melted LC2 promoter confirmed the bimodal FRET distribution with two sub-populations having mean E* values of 0.224±0.002 and 0.290±0.045 (**Fig. S5E**).

Consistent with the above equilibrium FRET values, 32% (N = 21) of real-time promoter binding trajectories in the presence of Myx inhibitor demonstrated the formation of initial RP_C_ complex (E*~0.2) and its subsequent isomerisation to E* ~0.3 state (**Fig. S5F**) while the remaining 68% (N = 45) of the nascent RNAP–promoter complexes maintained the E*~0.2 state for the entire duration of the trajectory (**Fig. S5G**). The increasing trend in observed FRET values as the RNAP–promoter complexes react towards RP_O_ is consistent with structural modelling data; the distance between the −15 and +15 labels decreases from 98 Å in the RP_C_ complex, to 87 Å in the Cor-stabilised RNAP-promoter intermediate, and further to 66 Å in the RP_O_ complex (**Fig. S5H**). We thus suggest that the RP_i_ intermediate, which we detected in a subpopulation of the RP_O_ formation FRET trajectories, resembles the Myx/Cor-stabilised RNAP-promoter complex previously characterised by cryo-EM (9) with respect to the downstream DNA residing in the RNAP cleft and the exclusion of the template DNA from the active site (**Fig. S5H**). Consistent with these complexes resembling a real on-pathway intermediate, Boyaci *et al.* (9) identified by cryo-EM a very similar RNAP–promoter structure (named as RP_i2_ in *ref.* (9)) also in the absence of Cor inhibitor (**Fig. S5H**).

### Transcription bubble opening leads to static and dynamic RP_O_ complexes

We next analysed the transcription bubble behaviour immediately *after* initial RP_O_ complex formation; our observation span for these measurements was 1.3–22 s (median = 4 s, N = 119). HMM analysis of the trajectories revealed two RP_O_ complex sub-populations: a “static” (or “stable”) sub-population (73% of the nascent RP_O_ complexes; exact 95% binomial CI: 64–81%), where the transcription bubble remained open for the entire duration of the trajectory (reflected by an E*~0.45 state; **Fig. 4A**); and a dynamic sub-population (27%, CI: 19–36%), where the transcription bubble fluctuates rapidly between the open state (E*~0.45) and state(s) characterised by lower FRET (**Fig. 4B**). The dynamic RP_O_ complexes do not appear to convert to a static RP_O_ within our observation span, suggesting that the dynamic RP_O_–like complex is not an on-pathway intermediate which eventually isomerises to form the stable RP_O_ complex. This conclusion, which invokes the formation of static and dynamic RP_O_ complexes on parallel pathways, is also supported by the presence of a similar distribution of static and dynamic RP_O_ complexes in samples of preformed, heparin-challenged RNAP–promoter complexes (see next section of the manuscript and ref. (43)). Notably, both the 2-state (i.e., showing no RP_i_ intermediate; **Fig. 2B**) and 3-state (**Fig. 3A**) bubble-opening mechanisms produced both static and dynamic RP_O_ complexes (**Fig. 4C**), suggesting that the presence of the RP_i_ intermediate does not dictate the subsequent stability of the bubble.

**Figure 4.**
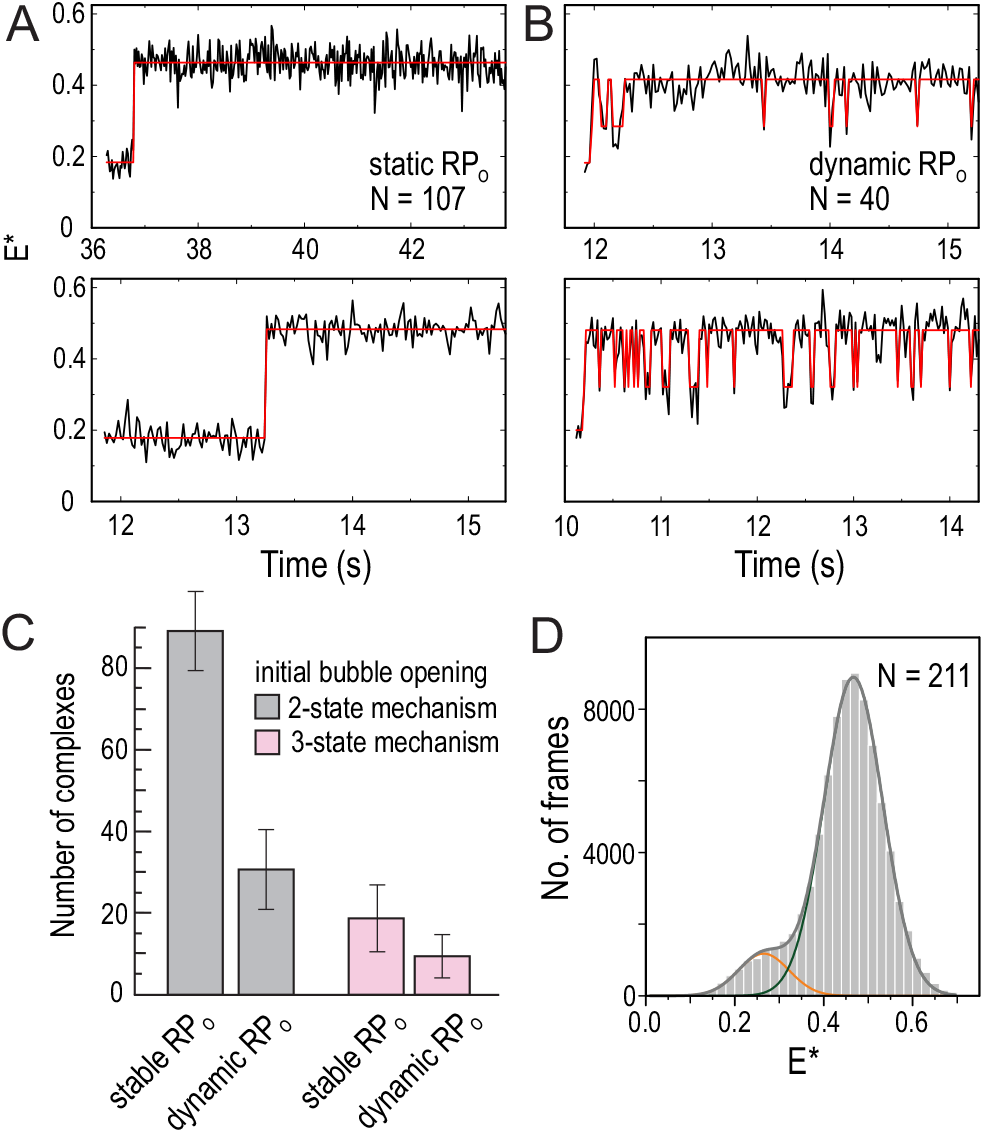
Parallel formation of static and dynamic RP_O_’s. (A) Example trajectories demonstrating the formation of static RP_O_. (B) Example trajectories demonstrating the formation of dynamic RP_O_. The static population represents 73% (N=107) of the nascent RP_O_ complexes, whereas the dynamic population represents 27% of the complexes (N=40). (C) The abundance of static and dynamic RP_O_’s is shown separately following the initial bubble opening either via 2-state (grey bars) or 3-state (pink) mechanisms. Exact 95% binomial confidence intervals are shown. (D) E* histogram of dynamic RP_O_’s. The complexes were imaged ~5 min after their initial formation on the coverslip surface (N = 211 molecules). The E* distributions were fit using Equation 2.

To evaluate the bubble conformations accessed by the dynamic RP_O_ complexes, we analysed the E* distribution of these complexes, which was fit well by two Gaussian functions with mean E* values of 0.265±0.004 and 0.467±0.001 (**Fig. 4D**). In contrast, the E* histogram of the static RP_O_ showed only a single distribution centred at E* of 0.448±0.001 (**Fig. 3B**). These values suggest that the dynamic RP_O_ complexes do not sample either the RP_C_ (E*~0.20) or the on-pathway intermediate RP_i_ (E*~0.32) states. Instead, it is more likely that the dynamic RP_O_ complexes sample an off-pathway state, which is characterised by E*~0.27 and which we coin as RP_ISO_.

### The role of RNAP–promoter upstream interactions in the RP_O_ pathway selection

We next evaluated how RNAP αCTD–promoter upstream sequence interactions contribute to the relative formation of stable and dynamic RP_O_ complexes and the kinetic parameters of the transcription bubble dynamics. To this end, we prepared RP_O_ complexes at 37°C using either a short or a long dsLC2, challenged them with heparin, i.e., a DNA competitor, and immobilised them on the coverslip surface for smFRET analysis. In this protocol, RP_O_ complexes also form on the short dsLC2 promoter fragments, allowing direct comparison to the long dsLC2 promoter (**Fig. S6A**). HMM-based classification of the trajectories indicated that the dynamic RP_O_ complexes were 1.7-fold more prevalent (25±3.6% vs 15±2.7% of all complexes; mean and SD of three independent experiments) on the short promoter compared to the long promoter (**Fig. S6B**). A two-sample *T*-test (*p*=0.035) confirmed that the observed difference in the relative number of dynamic complexes on the two promoters is statistically significant. Further, the observation span for the measurements was 2.0–25 s (median 8.0 s, N = 348) on the long promoter and 1.4–25 s (median 6.9 s, N = 435) on the short promoter, indicating that the higher prevalence of dynamic RP_O_ complexes on the short promoter is not explained simply by longer trajectories that accumulate more state transitions. The mean dwell-time of the dynamic complexes in the RP_ISO_ (85–90 ms, **Fig. S6C,E**) and RP_O_ (560–660 ms, **Fig. S6D,F**) states were similar on both promoters (**Fig. S6G**). Our results suggest that the αCTD–promoter interactions steer the RP_O_ pathway selection towards the static RP_O_ complex; however, the effect is moderate, and significant number of dynamic RP_O_ complexes form also on the long promoter. The similarity in the timescales of transcription bubble dynamics on the short and long promoters also indicates that the bubble isomerisation rates are independent of the status of the αCTD–promoter upstream sequence interactions.

### The role of RNAP cleft loops in the RP_O_ stabilisation

To interrogate protein structural elements contributing to the RP_O_ stability, we deleted the β gate loop (ΔGL), β’ rudder loop (ΔRL) or β’ lid loop (ΔLL) from the RNAP and determined the effects of these deletions on the transcription bubble dynamics using preformed RP_O_ complexes. Structurally, GL mediates the RNAP β-pincer interaction with the RNAP β‘-clamp, thus forming a barrier for the DNA entry and exit from the RNAP cleft (**Fig. 5A**) (9)(10); RL locates between the tDNA and ntDNA strands in the RP_O_; and LL locates adjacent to the RL and is able to interact with the tDNA around base −6 in the RP_O_.

**Figure 5.**
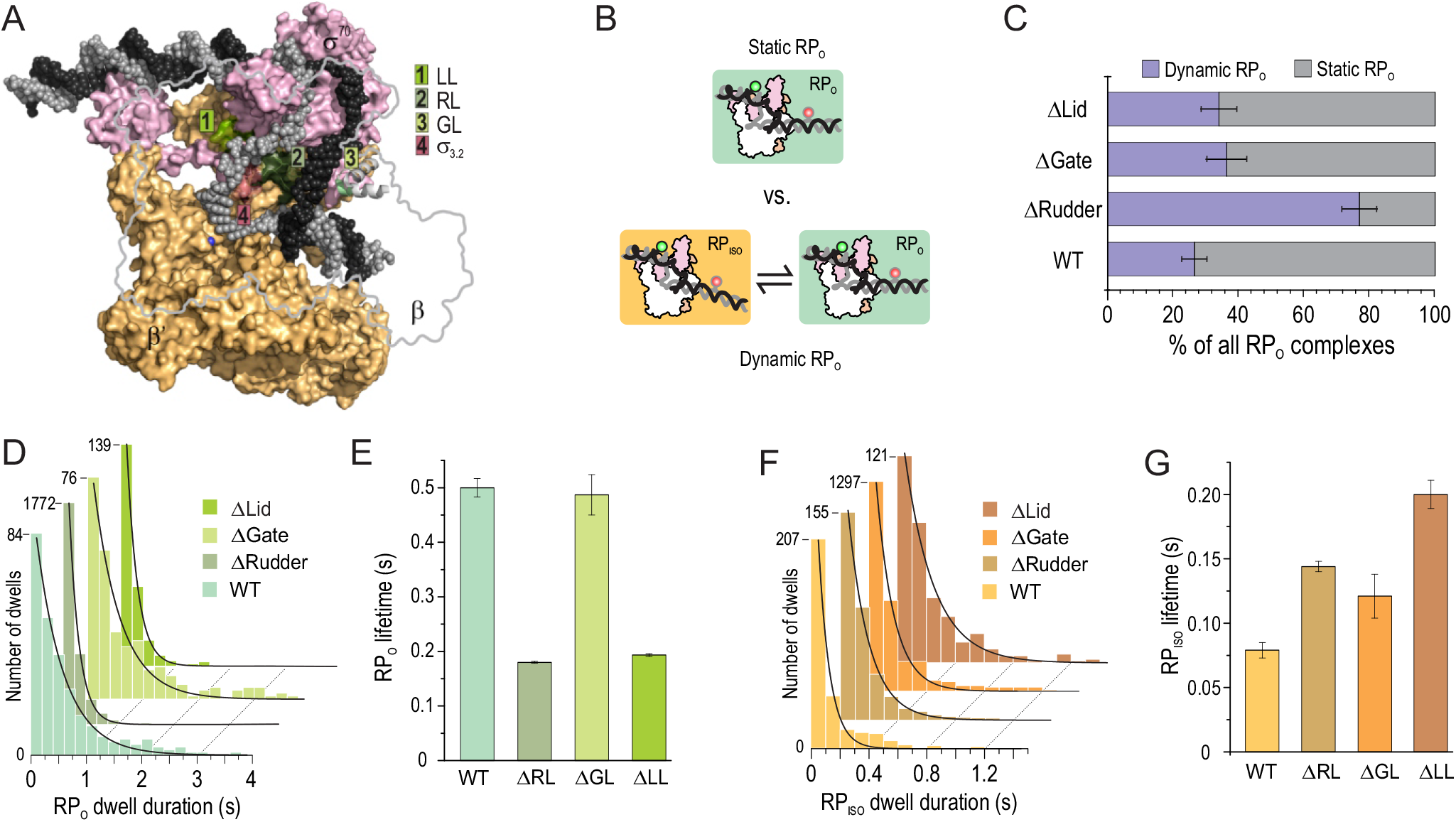
Effect of RNAP cleft loop deletions on the reaction pathway branching and transcription bubble kinetics. (A) The position of deleted lid loop (LL), rudder loop (RL) and gate loop (GL) are shown in the cryo-EM model of *E.coli* RP_O_ (PDB: 6psw, (10)). α, β and ω RNAP subunits and TraR transcription factor are omitted for clarity. ntDNA and tDNA are shown in dark and light grey, respectively. Blue sphere is the active site Mg^2+^ ion. (B) The RP_O_ complexes were classified as static or dynamic based on the 2-state HMM fit of the FRET trajectories. (C) The relative amounts of static and dynamic RP_O_. Error bar: exact 95% binomial CI. WT, N=212 molecules; ΔRL, N=206; ΔGL, N=136; ΔLL, N=115. (D) Dwell time distributions of the RP_O_ state within the dynamic RP_O_ population were fit to mono-exponential equation. (E) The lifetime of RP_O_ state was obtained from the dwell distributions in panel D. Error bars are 1 SE extracted from the fit. (F) Dwell time distributions of the RP_ISO_ state within the dynamic RP_O_ population were fit to mono-exponential decay equation. (G) The lifetime of RP_ISO_ state was obtained from the dwell distributions in panel F. Error bars are 1 SE extracted from the fit.

We used 2-state HMM of FRET trajectories to classify RP_O_ complexes into static (i.e., no bubble dynamics) and dynamic (**Fig. 5B**) and found that all deletions shifted the balance of RP_O_ formation towards the direction of dynamic RP_O_ complexes. The effects of ΔGL and ΔLL were moderate, as they increased the fraction of dynamic RP_O_ complexes from the wild-type (WT) RNAP level by 1.3–1.4-fold, i.e., from 26% (exact 95% binomial CI: 22–31%) in WT to 34% (CI: 29–40%) in ΔLL and 36% (CI: 30– 43%) in ΔGL (**Fig. 5C**). However, the ΔRL effect was much stronger, 2.9–fold, making most RP_O_ complexes, i.e., 77% (CI: 72–83%), dynamic. Kinetic analysis of the dynamic RP_O_’s indicated that ΔRL and ΔLL shortened the lifetime of the open bubble state by 2.8–fold, whereas ΔGL had no effect (**Fig. 5D,E**). In contrast, all three mutations increased the lifetime of the RP_iSO_ state. Specifically, the RP_iSO_ lifetime increased by 2.5-, 1.8- and 1.5-fold in the case of ΔLL, ΔRL and ΔGL mutant RNAPs, respectively (**Fig. 5F,G**). Notably, the median observation span for these measurements was 4.26 s, 4.84 s, 5.20 s and 6.00 s in the case of WT, LL, GL and RL, respectively. The variation in the observation span, however, does not significantly contribute to the classification of the molecules to the static and dynamic RP_O_ classes or kinetic parameters because, even in the combination of shortest trajectories (median 4.26 s) and most stable state, i.e., wild-type RP_O_ complex (lifetime 0.5 s), the probability that RP_O_ → RP_ISO_ transition does not take place within the trajectory is extremely small (only 0.0002). Collectively, our data indicate that the GL, RL and LL domains in the RNAP contribute to the RP_O_ pathway branching (by changing the balance between static and dynamic RP_O_ complexes) and affect the rates of the transcription bubble conformation changes.

## Discussion

In this work, we establish the ability to look at the earliest stages of transcription initiation in real-time and at the single-molecule level. This unique capability bypasses the need to synchronise complexes and offers unprecedented access to co-existing reaction pathways and transient intermediates; as a result, we gained new mechanistic insight about the paths and intermediates used by RNA polymerase to form σ^70^-dependent RP_O_ complexes on a *lac* promoter derivative. Our work also provides further insight on the dynamics and heterogeneity of open complexes and their determinants, as well as their potential biological role in gene regulation.

### Open complex formation may proceed via more than one path

Our data indicate that the RP_C_ → RP_O_ isomerisation step involves **mechanistic branching**. In most cases (80–86% of molecules), the isomerisation occurred in one apparent step without observable intermediate(s) within our 20-ms temporal resolution, suggesting that bubble initiation was followed by rapid bubble progression, downstream DNA loading to the RNAP cleft, and template DNA loading to the active site cleft. However, the remaining molecular trajectories (14-20%) displayed an **intermediate state (RP_i_**) between the RP_C_ and RP_O_. Given the RP_i_ lifetime and our temporal resolution, we excluded the possibility that all the trajectories without an identified RP_i_ are due to its short lifetime; instead, our data support the presence of parallel paths for RP_O_ formation, or at least, the presence of RNAP populations with different propensity to form an open complex.

### Possible structure of RP_i_

Based on the observed FRET efficiencies, the intermediate may have structural similarities to an RNAP-promoter complex stabilised by an antibiotic targeting RNAP. Recent cryo-EM data of *Mycobacterium tuberculosis* RNAP showed that corallopyronin A (a Myx analogue) stabilises a partially melted transcription bubble (region −11/−4) (9), consistent with earlier biochemical footprinting data (45). The same cryo-EM work showed that a similar conformation was present in the absence of Myx, raising the possibility that the observed conformation may represent an intermediate on the RP_O_ pathway (9). Our real-time trajectories using *E. coli* RNAP clearly show the presence of an intermediate (RP_i_) that has a structural signature similar to that expected by a partially melted bubble. Another possibility for the structure of the RP_i_ is an intermediate where all the melting has been completed, but the template DNA has not been loaded to the active site cleft; such a structure is supported by results showing that the presence of Myx does not prevent full opening of DNA, as sensed by fluorescence enhancement of a Cy3 probe introduced at position +2 (46); such enhancement is expected when the bubble opening has been complete. However, at least on *lac* promoter, the intermediate is kinetically significant only in a small fraction of RP_O_ formation events.

### Possible sources for heterogeneity in open complex formation pathways

Since the **RP_i_** is detectable only in a subset of trajectories, it is conceivable that, for those trajectories, a structural module of the RNAP delays downstream bubble melting and/or tDNA loading to the active site cleft. One candidate for such a structural module is the RNAP **βʹ switch-2 segment**. Based on mutational analysis, structural studies, and observed Myx effects, it has been established that the switch-2 region can adopt two different conformations (45). The conformation dominant in the absence of Myx is compatible with template loading to the active site; in contrast, the alternative conformation (which is stabilised by Myx and specific mutations in the switch-2) blocks template loading to the active site. If the switch-2 was at the moment of DNA bubble initiation in the blocking conformation in a small fraction of RNAPs, the loading of template DNA to the active site cleft would be delayed by the necessary switch-2 refolding. This hypothesis also predicts that the kinetics of the switch-2 conformational change (from the blocking conformation to the permissive conformation) controls the rate of formation of RP_O_ in a population of RNAPs.

The RP_i_ heterogeneity may also result from alternative **promoter discriminator** (region −6/+1) conformations in different RNAP molecules. This hypothesis is supported by the previous finding that G^−6^G^−5^G^−4^ and C^−6^C^−5^C/T^−4^ motifs in the ntDNA stabilised *in crystallo* two distinct discriminator conformations and imposed in solution one base-pair difference in the predominant transcription start site (47). GTG motif, which is found in our promoter, had transcription start site statistics halfway between the GGG and CCC/T motifs, consistent with the assumption that a promoter with this motif can readily adopt either of the two discriminator conformations.

### RP_O_ complexes appear to have different stability immediately after their formation

A longstanding question in the transcription field is whether all RP_O_ on a given promoter are the same or differ in their structural and functional properties (22)(23)(24)(25). Our results show that indeed there is another layer of heterogeneity as indicated by the **differing stability of the transcription bubble,** even immediately after RP_O_ formation (as judged by the appearance of the E*~0.45 state). Most RP_O_ complexes can keep the bubble open for at least several seconds; however, a more dynamic RP_O_ subpopulation samples different bubble states in the millisecond timescale. The stable or dynamic RP_O_ mode was set before or during the first-time opening of the bubble and the modes did not interconvert within our observation span (~8 seconds); this observation rules out the possibility that the dynamic RP_O_ were indeed intermediates on the linear pathway leading to the stable RP_O_ complexes. This new insight aligns well with our previous observation of stable and dynamic RP_O_ molecules within the population of pre-formed RP_O_ complexes (43), while allowing further mechanistic insight by linking the formation of stable and dynamic complexes on the existence of a branched RP_O_ pathway and the sampling of a short-lived off-pathway state (RP_ISO_) by the dynamic RP_O_’s.

The analysis of **mutant RNAPs** suggest that the main difference between the dynamic and stable RP_O_ complexes arises from the RNAP interaction with the single-stranded template DNA in the active site cleft. The deletion of rudder loop, which presses against the template DNA positions −7 to −9, tripled the amount of dynamic RP_O_ (from 26% to 77%; **Fig. 5C**) and decreased 3-fold the open bubble lifetime in the dynamic RP_O_ complexes (**Fig. 5E**). The deletion of lid loop, which interacts with the template DNA base −6, had a similar effect on the open bubble lifetime. We previously found that deletion of the σ^70^ 3.2 region (the σ “finger”, which interacts with the template DNA strand from bases −3 to −6), destabilised the RP_O_ (43). These interactions with the template DNA form late in the RP_O_ mechanism, i.e., when the bubble forms fully and the template DNA strand loads to the active site cleft. Our data suggest that these final interactions form by two alternative ways generating “tight” and “loose” template DNA binding modes: the tight binding mode gives rise to the stable RP_O_ complexes, and the loose binding mode gives rise to the dynamic RP_O_ complexes.

The presence of such dramatic differences in RP_O_ stability may have functional consequences, and may be related to reports of non-uniform RP_O_ function. Specifically, a subset of RP_O_ complexes (on many promoters) appear to be locked in an abortive initiation mode, in which they repetitively synthesise short RNA products (<12-mer, with the exact sequence depending on the specific promoter), whereas another RP_O_ subset escape the promoter efficiently and synthesise full-length RNA products (22)(23)(24)(25). The failure of promoter escape may also result from long-lived backtracking and arrest of initially transcribing RNAPs (28)(29). The RP_O_’s apparently locked in the abortive mode are also referred as “moribund” complexes, and they apparently have a role in transcription regulation in the cell (27). Mechanistically, the dynamic RP_O_ could be candidates to form such moribund complexes, since unstable template DNA binding to the active site is likely to enhance the dissociation probability of short RNAs, leading to abortive initiation. Consistent with this reasoning, the Δ3.2 σ^70^ mutant (which increase RP_O_ dynamics substantially) released 4-7-mer RNAs more efficiently compared to the WT (24).

### Role of the promoter upstream interactions on the RP_O_ formation and stability

The **RNAP** α**CTDs** interact with the promoter upstream sequences either by specifically recognising the promoter UP element (18) or via sequence-independent interactions (14)(19); both interactions are important for RP_O_ formation. We found that the upstream part of the *lacCONS+2* promoter (from −40 to −89; **Fig. S2A-B**), which does not contain a full UP element but is similar to the distal UP subsite (48), facilitates transcription bubble melting in the context of fully double-stranded promoters. In fact, the short double-stranded promoter (which lacks αCTDs interactions) failed to form RP_O_ under our experimental conditions, which involve measurements at room temperature. On the other hand, if the requirement for the DNA melting nucleation was bypassed (e.g., by using a pre-melted promoter), the αCTD interactions with upstream sequence no longer increased the rate of RP_O_ formation (**Fig. 2D,E**). This finding is consistent with previous biochemical studies showing that the αCTD interactions with the UP element enhance both the initial promoter binding and subsequent isomerisation to competitor-resistant conformation (14)(19).

We also found that the presence of upstream sequence interactions did not substantially change the ratio of initial bubble opening events that occur in one step or in two steps (i.e., via the RP_i_), and did not significantly change the rates of transcription bubble dynamics in the pre-formed RP_O_. However, the dynamic RP_O_ complexes formed more often on the short promoter in comparison to the long promoter (25% vs. 16%), suggesting that the αCTD–promoter interactions, instead of being fully decoupled from mechanistic steps occurring after the bubble nucleation, have a role in the late steps of RP_O_ pathway branching; the exact mechanism of such modulation is unclear, but it may involve bending of the upstream sequence on the RNAP surface (17) and subsequent interactions that affect RNAP conformation dynamics in a way that it influences bubble dynamics.

### A working model for the RP_O_ formation mechanism

We summarise our key findings in the context of the RP_O_ formation mechanism in **Fig. 6** (2). The process begins with the RP_C_ complex formation as the RNAP holoenzyme binds to the promoter and establishes interactions with the −35 element, −10 element and upstream sequences. Interaction of αCTDs with upstream sequences stimulate RP_O_ formation by bending the upstream DNA around the RNAP (12)(13)(14)(15)(16)(17) and coupling it energetically with bubble formation.

**Figure 6.**
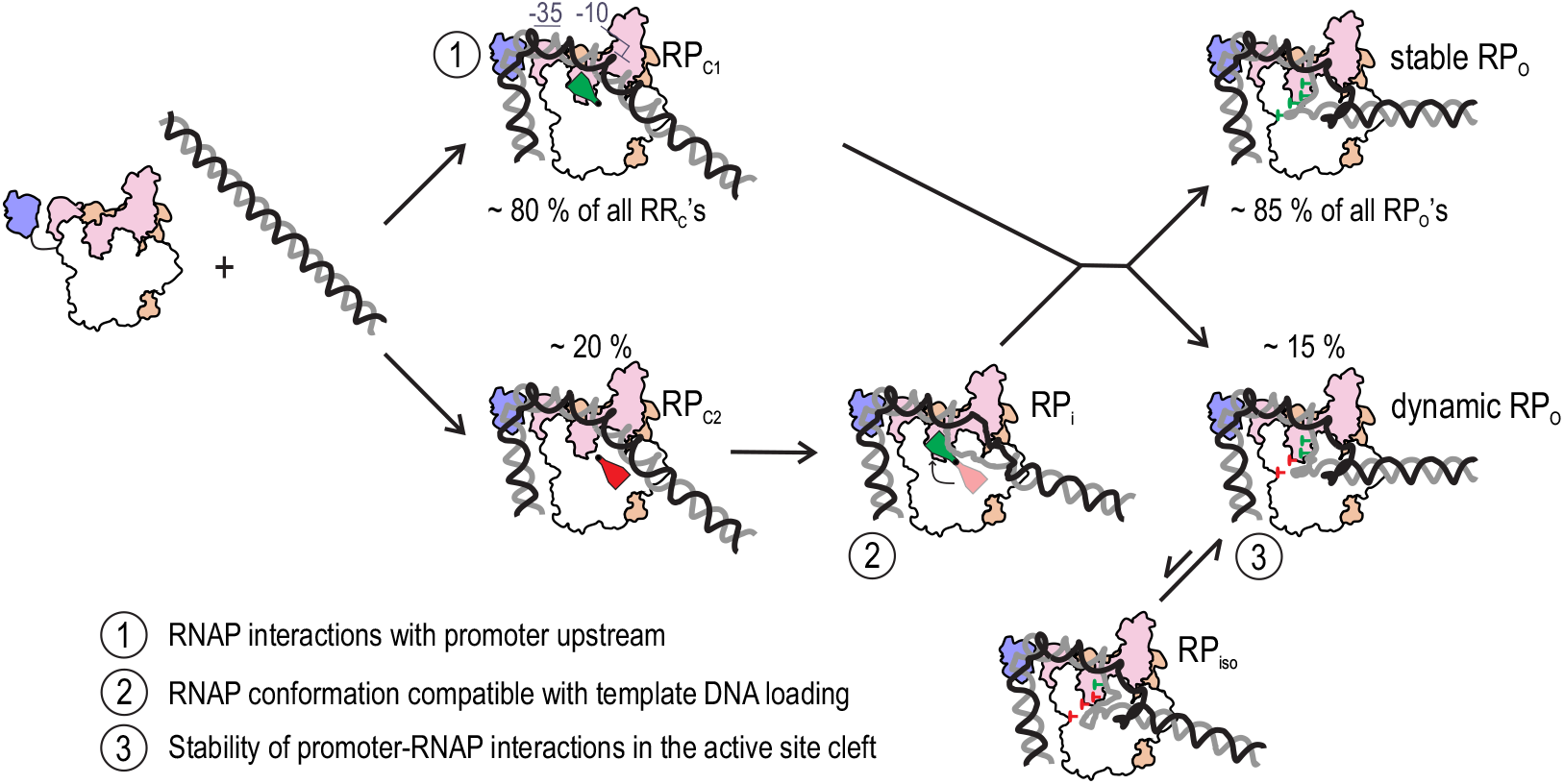
A working model for the RP_O_ complex formation mechanism. The reaction pathway from the promoter binding to the RP_O_ complex is branched in two separate steps. The first branching is hypothesised to depend on a mobile RNAP element, which can be either in an active or inactive conformation (green and red flaps, respectively). The inactive conformation blocks the loading of the tDNA strand into the active site cleft, resulting the formation of intermediate RP_i_. The isomerisation of the mobile element to the active conformation clears the block and allows progress from RP_i_ to RP_O_. The second branching is related to the stability of the RNAP–template DNA interaction in the active site cleft. In ~15% of the RP_O_ complexes, the interaction is weak, allowing continuous dynamic movement of the template DNA and thus the downstream DNA. Because stable and dynamics RP_O_ complexes formed both from RP_C_1 and RP_i_, we assume that these pathways merge before the next branching step leading to the stable and dynamics RP_O_’s. Green vs red pins depict tight and loose interactions between the tDNA and the RNAP, respectively. The numeration (1, 2 and 3) indicates the key steps in the mechanism that define the rate and efficiency of RP_O_ complex formation and that may be modulated by the promoter sequence and transcription regulators.

The initial nucleated bubble expands via two different mechanistic paths: in the first path (most common for our *lac* promoter derivative), the RNAP melts the entire bubble and loads the template DNA strand to the active site cleft in one apparent step without detectable intermediates; the second path, however, involves a short-lived intermediate, RP_i_, which features partial bubble melting or incomplete template loading to the active site cleft. We hypothesise that the intermediate appears when a mobile element of the RNAP, e.g., the switch-2 module, is initially in a conformation incompatible with template loading to the active site cleft or if an alternative promoter discriminator conformation delays the full bubble melting in a subset of RNAP–promoter complexes. Template DNA loading to the active site cleft leads to the tight-binding and loose-binding states, which do not readily interconvert. Because stable and dynamics RP_O_ complexes formed with similar probability both directly from RP_C_ and via RP_i_, we assume that these pathways merge before the branching to the stable and dynamics RP_O_’s takes place at the template DNA loading step (**Fig. 6**). The tight template DNA binding mode keeps transcription bubble open whereas the loose-binding features dynamic movement of the template DNA. Template DNA interactions with the RNAP rudder loop and σ finger are part of the key determinants of tight binding mode.

Within this mechanistic framework, transcription initiation can be modulated by transcription factors or small-molecule regulators that change RNAP interactions with upstream sequences since such interactions strongly affect promoter binding (14)(19) and DNA melting kinetics. The pathway branching to static and dynamic RP_O_ may be the underlying cause of complexes being locked in abortive initiation (22)(23)(24)(25). The inherent mechanistic heterogeneity of the RP_O_ formation pathway may amplify the effects of transcription regulators and promoter sequence by converting relatively modest perturbations into large changes in the formation rate, stability and functional properties of the RP_O_ complexes. For example, GreB transcription factor was previously found to block the RP_O_ pathway to a step after the RP_C_ formation (49) whereas copper efflux regulator CueR modulated the formation probability of active and inactive RNAP–promoter complexes (50).

## Abbreviations

ALEX: Alternating laser excitation
αCTD: C-terminal domain of RNA polymerase α-subunit
CI: confidence interval
Cor: corallopyronin A
cryo-EM: cryo-electron microscopy
E*: apparent FRET efficiency
FRET: fluorescence energy transfer
GL: gate loop in the RNAP β’subunit
HMM: Hidden Markov modelling
LL: lid loop in the RNAP β’ subunit
Myx: myxopyronin B
RL: rudder loop in the RNAP β’ subunit
RNAP: RNA polymerase
RP_C_: RNAP–promoter closed complex
RP_i_: RNAP–promoter intermediate complex
RP_O_: RNAP–promoter open complex
SD: standard deviation
SE: standard error

## Data availability

All our time-trace data and the HMM software we used for their analysis will be available to any interested party upon request.

## Funding

This work was supported by Academy of Finland [grant numbers 307775, 314100, 335377 to A.M.M]; Instrumentarium Science Foundation [grant to A.M.M.]; Finnish Cultural Foundation [grants to A.M.M. and O.P.]; Wellcome Trust grant [grant number 110164/Z/15/Z to A.N.K.]; and UK Biotechnology and Biological Sciences Research Council [grant number BB/H01795X/1 to A.N.K].

## Acknowledgements

We thank Richard E. Ebright for providing myxopyronin B. We are grateful to Matti Turtola, Thadée Grocholski and Henri Malmi for constructing plasmids. We thank Abhishek Mazumder for insightful discussions and critical reading of the manuscript. We thank the Cell Imaging and Cytometry core at Turku Bioscience Centre, which is supported by Biocenter Finland, for the access to the microscopes.

## Notes

### Competing Interest Statement

The authors have declared no competing interest.

