## Supplementary information for "The formation of the bacterial RNA polymerase–promoter open complex involves a branched pathway"

Supplementary Figure captions

Figure S1. Promoter sequences used in the study. (A) The DNA sequences and fluorophore positions of the long and short fully double-stranded lacCONS+2 promoters used in this study are shown. Non-template DNA (ntDNA) strand is in black, template DNA (tDNA) strand is in grey. Donor fluorophore Cy3B (green sphere) is attached to the -15 ntDNA thymine residue. Acceptor fluorophore ATTO647N (A647N, red sphere) is attached to the +15 tDNA thymine residue. (B) The long and short pre-melted versions of the lacCONS+2 promoters were created by changing the template DNA sequence between the positions -10 and -4 as indicated in blue. (C) The short pre-melted version of the lacCONS+2 promoter with acceptor ATTO647N label attached to the +20 tDNA thymine residue. This promoter was specifically used to demonstrate the transcriptional activity of formed RNAP–promoter complexes in Fig. S3. (D) Differences in the ntDNA sequence of the used lacCONS+2 and native E. coli lac promoter are shown in orange. The substitutions in the CAP binding site in promoter upstream extension, the -35 element and the -10 element as well as the single basepair deletion in the -35/-10 spacer change these promoter elements to their consensus versions (1) (2).

Figure S2. Specificity of promoter DNA binding to the surface-immobilised RNAP holoenzyme. (A) E. coli RNAP–$\sigma^{70}$ holoenzyme was immobilised on the PEGylated coverslip. The immobilisation chemistry included PEG/biotin-PEG layer on the coverslip surface, neutravidin and biotinylated anti-His-tag-antibody. Short pre-melted lacCONS+2 promoter (length -39/+25) labelled with Cy3B donor fluorophore at non-template DNA position -15 and ATTO647N acceptor fluorophore at template DNA position +15 was added to the reaction buffer and let to bind 5 min. The coverslip was then imaged with Nanoimager S microscope (ONI, Oxford, UK) using the total internal reflection mode. The objective magnification was 100X and numerical aperture 1.4. Cy3B fluorophore was imaged on the DD channel using 561 nm laser and ATTO647N fluorophore was imaged on the AA channel using 640 nm laser. The frame time per channel was 20 ms. The pixels of the raw images were 2 x 2 binned (sum) using Fiji software (3); a single frame on each channel is shown. (B) Negative control experiment was performed as in panel A with the exception that the RNAP holoenzyme was omitted from the coverslip surface. (C) The E* histograms demonstrate the formation of $RP_O$ (E~0.45) on the coverslip surface (grey, N=291 molecules). $RP_O$, i.e. the E~0.45 promoter state, disappears after 2 min incubation with 1 mM NTP's (orange, N=81) as the RNAP starts RNA synthesis and escapes the promoter that leads to promoter DNA conformation with low FRET efficiency. The E* values were extracted from each frame (20 ms) of the recorded movies. The promoter was long fully double-stranded lacCONS+2 with Cy3B at -15 ntDNA and ATTO647N at +15 tDNA. (D) The E* histogram of a biotinylated protein-free promoter (N=61). The data were collected as in panel C.

Figure S3. Transcription activity of surface-formed RNAP–promoter complexes. (A) The activity of the RNAP–promoter complexes was monitored using short pre-melted lacCONS+2 promoter (length -39/+25) labelled with Cy3B fluorophore at non-template DNA position -15 and ATTO647N fluorophore at template DNA position +20. This previously developed labelling scheme creates a FRET ruler, which shows characteristic low FRET for the $RP_O$ (and inactive RNAP–DNA complexes), intermediate FRET for initially transcribing complexes (ITCs) containing 4–7-mer RNA and high FRET for the ITCs containing >7-mer RNA, respectively. After RNAP escape from the promoter and the formation of transcription elongation complex (TEC) FRET returns to low level (4). (B) Schematic FRET trajectory demonstrating the formation of RNAP–promoter complex at the beginning of FRET trace and subsequent increase in the FRET as the $RP_O$ complex forms and commences RNA synthesis. The FRET signatures highlighted with turquoise and purple demonstrate the events of abortive initiation and promoter escape, respectively. (C) Four experimental trajectories demonstrating transcription activity shortly after the formation of the RNAP–promoter complex. The RNAP was immobilised to the coverslip surface as described in Fig. S2. The imaging buffer contained 1 mM ATP, GTP, CTP and UTP; at the ~10 s time-point, 1 nM promoter DNA was added to the sample well. The frame time of the recordings was 200 ms with 100 ms ALEX excitation by green and red laser, respectively. Approximately 51% of all molecules (N=80 total molecules) showed FRET signatures indicating transcriptional activity. (D) Two experimental trajectories demonstrating the formation of inactive RNAP–promoter complexes. Approximately 49% of molecules showed FRET signatures of inactive RNAP–promoter complexes, i.e., static E* ~0.1–0.2.

Figure S4. The formation efficiency of $RP_O$ complex on different lacCONS+2 promoters. The E* histograms reflect the formation efficiency of the open transcription bubble and $RP_O$ complex (E*~0.45 peak) on (A) the short double-stranded LC2 promoter, (B) long dsLC2 promoter, (C) short pre-melted LC2 promoter and (D) long pre-melted LC2 promoter, respectively. RNAP–promoter complexes were recorded ~5 min after their initial formation on the coverslip surface. The FRET values were extracted from each frame (20 ms) of the recorded movies. The promoters were labelled with Cy3B at ntDNA position -15 and ATTO647N at tDNA position +15. Data statistics: short double-stranded LC2 promoter N=90 molecules and 35 400 frames; long double-stranded LC2 promoter N=675 molecules and 255 000 frames; short pre-melted LC2 promoter N=388 molecules and 99 600 frames; long pre-melted LC2 promoter N=291 molecules and 89 500 frames. Blue and green rectangles indicate expected E* values for the closed and open transcription bubble conformations, respectively. The E* ~0.2 species in the blue rectangle contain also unspecific RNAP–promoter complexes.

Figure S5. Characterisation of RNAP-promoter complexes formed in the presence of myxopyronin B inhibitor. (A) Preformed RNAP–promoter complexes were classified into three classes based on their FRET behavior. Class 1 molecules comprised of RNAP–promoter–Myx inhibition complexes, which are characterised by E* ~0.3 state. (B) A subset of RNAP–promoter–Myx inhibited complexes sampled a short-lived higher E* state. Typically this higher E* state lasted only 1–2 frames (20–40 ms). (C) Class 2 molecules are characterised by stable E* ~0.2 state and probably constitute of non-productive RNAP-promoter complexes. (D) Class 3 molecules are $RP_O$ complexes, which are identified based on their long-lived E* ~0.45 state. These few identified $RP_O$ complexes probably formed after the dissociation of Myx inhibitor. The promoter in panels A–D was long dsLC2. (E) FRET efficiency histogram of the RNAP–promoter complexes preformed in the presence of Myx on long pmLC2 promoter. The fit of E* distribution to Equation 2 identified mean E* values of 0.224±0.002 and 0.290±0.045. (F) Two example trajectories demonstrating the real-time formation of initial $RP_C$ complex (E*~0.2) and its subsequent isomerisation to E* ~0.3 state (N = 21). (G) Two example trajectories demonstrating the real-time formation of RNAP–promoter complex that remained in the E*~0.2 state for the entire duration of the trajectory (N = 45). The molecules in panels A–E were imaged ~5 min after the initial formation of the RNAP–promoter complexes on the coverslip surface. The molecules in panels F and G were imaged in real-time allowing the resolution of promoter binding and subsequent promoter conformation changes. The red solid line represents the Viterbi for a 2-state HMM. Myx concentration was 100 µM. (H) The location of Cy3B and ATTO647N fluorophores at the non-template DNA position -15 and template DNA +15, respectively, was modelled using FPS software (5) and indicated cryo-EM based RNAP structures. Noteworthy, corallopyronin was suggested to trap RNAP to a state with close similarity to a normal intermediate $RP_{i2}$ in the $RP_O$ formation pathway (6). The cryo-EM structure of $RP_C$ was obtained using E. coli proteins (7) whereas $RP_{Cor}$, $RP_{i2}$ and $RP_O$ structures were obtained using M. tuberculosis proteins (6). The distance between the average positions of modelled -15/+15 fluorophores in each RNAP state is shown. The protein elements, i.e., $\beta$' RNAP subunit (light orange) and $\sigma$ factor (pink), are shown as in the $RP_O$ cryo-EM model (6).

Figure S6. Effect of $\alpha$CTD–promoter interactions on the reaction pathway branching and the rates of transcription bubble dynamics. (A) Dynamic $RP_O$ complex samples $RP_{ISO}$ state while static $RP_O$ does not show any FRET transitions. (B) The formation frequency of stable and dynamic $RP_O$ complexes on short (-39/+25) and long (-89/+25) lacCONS+2 promoters. (C–F) The dwell time distribution of $RP_{ISO}$ (yellow) and $RP_O$ (green) states in the population of dynamic $RP_O$. The dwell time distributions were fit to mono-exponential equation. (G) The mean lifetime (t) of the $RP_{ISO}$ (yellow) or $RP_O$ (green) state. $RP_O$ complexes were prepared in solution at elevated temperature (37$^o$C), challenged with competitor (heparin) and immobilised on the coverslip surface for smFRET analysis at 22$^o$C.

### Figure S1

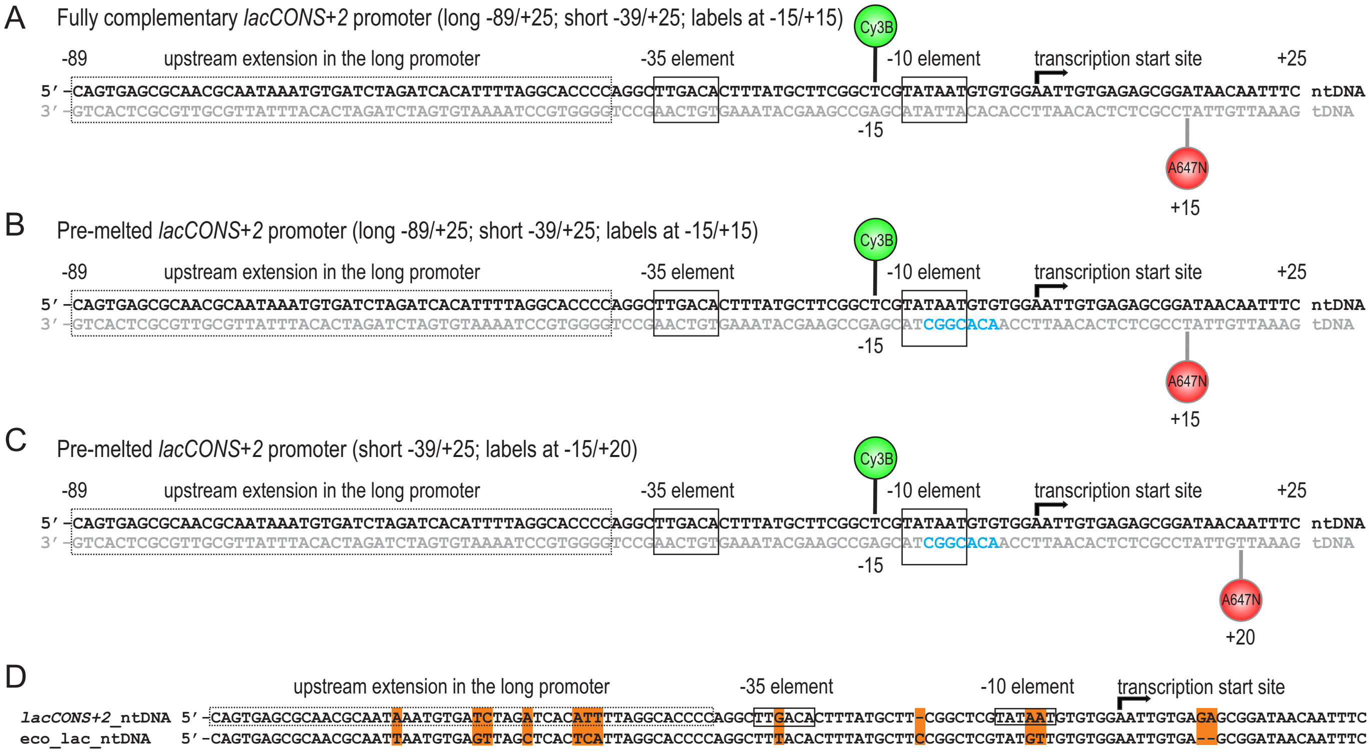
A
Fully complementary lacCONS+2 promoter (long -89/+25; short -39/+25; labels at -15/+15)
B
Pre-melted lacCONS+2 promoter (long -89/+25; short -39/+25; labels at -15/+15)
C
Pre-melted lacCONS+2 promoter (short -39/+25; labels at -15/+20)
-89
upstream extension in the long promoter
-35 element
-10 element
transcription start site
+25
Cy3B
A647N
-15
+15
+20
5′-CAGTGAGCGCAACGCAATAAATGTGATCTAGATCACATTTTAGGCACCCCAGGCTTGACACTTTATGCTTCGGCTCGTATAATGTGTGGAATTGTGAGAGCGGATAACAATTTC ntDNA
3′-GTCACTCGCGTTGCGTTATTTACACTAGATCTAGTGTAAAATCCGTGGGGTCCGAACTGTGAAATACGAAGCCGAGCATATTACACACCTTAACACTCTCGCCTATTGTTAAAG tDNA
3′-GTCACTCGCGTTGCGTTATTTACACTAGATCTAGTGTAAAATCCGTGGGGTCCGAACTGTGAAATACGAAGCCGAGCATCGGCACAACCTTAACACTCTCGCCTATTGTTAAAG tDNA
D
lacCONS+2_ntDNA 5′-CAGTGAGCGCAACGCAATAAATGTGATCTAGATCACATTTTAGGCACCCCAGGCTTGACACTTTATGCTT-CGGCTCGTATAATGTGTGGAATTGTGAGAGCGGATAACAATTTC
eco_lac_ntDNA 5′-CAGTGAGCGCAACGCAATTAATGTGAGTTAGCTCACTCATTAGGCACCCCAGGCTTTACACTTTATGCTTCCGGCTCGTATGTTGTGTGGAATTGTGA--GCGGATAACAATTTC

### Figure S2

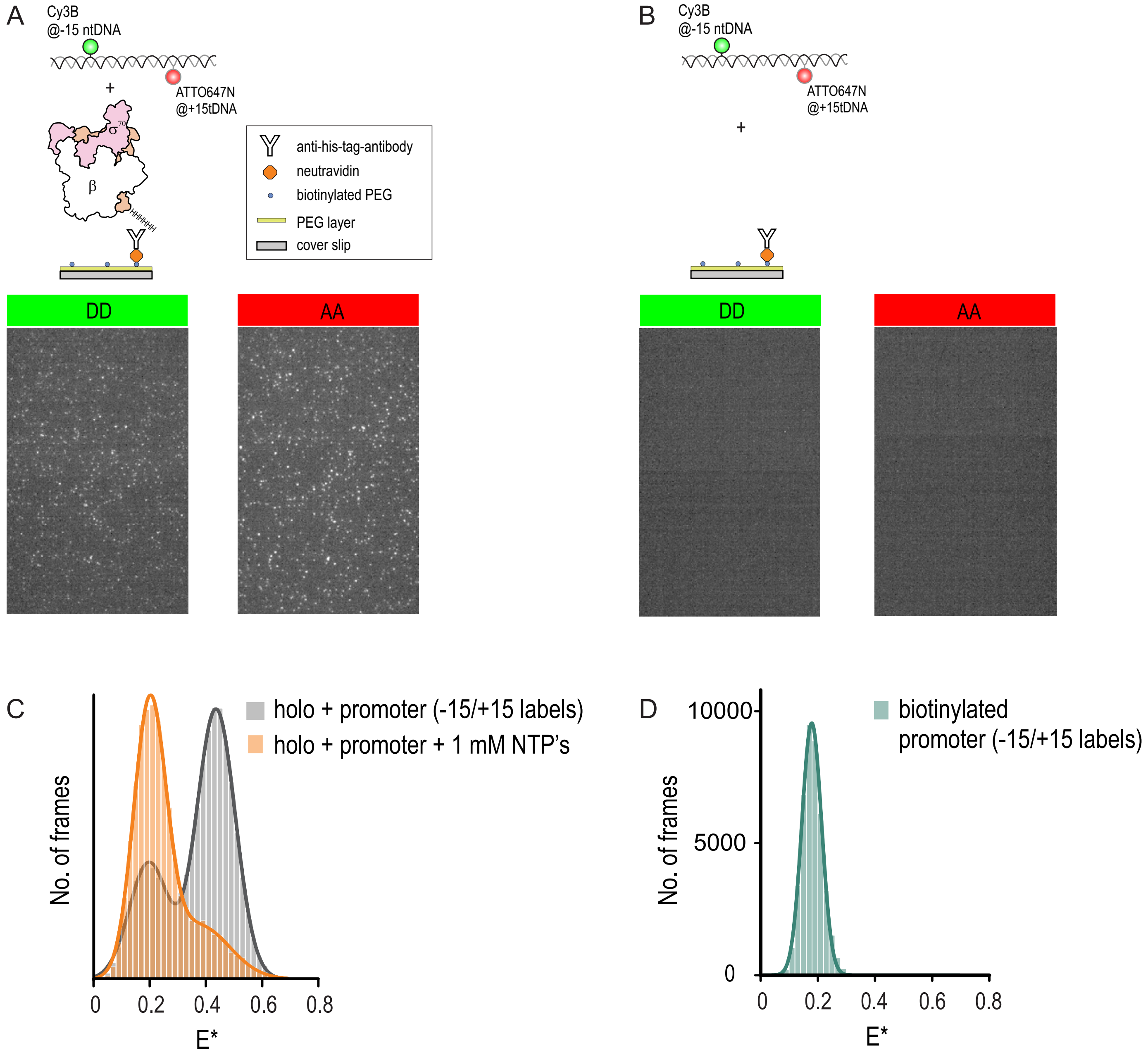
A
Cy3B
@-15 ntDNA
ATTO647N
@+15tDNA
+
σ
70
β
HHHHHH
anti-his-tag-antibody
neutravidin
biotinylated PEG
PEG layer
cover slip
DD
AA
B
Cy3B
@-15 ntDNA
ATTO647N
@+15tDNA
+
DD
AA
C
holo + promoter (-15/+15 labels)
holo + promoter + 1 mM NTP’s
No. of frames
0
0.2
0.4
0.6
0.8
E*
D
biotinylated
promoter (-15/+15 labels)
10000
5000
0
No. of frames
E*

### Figure S3

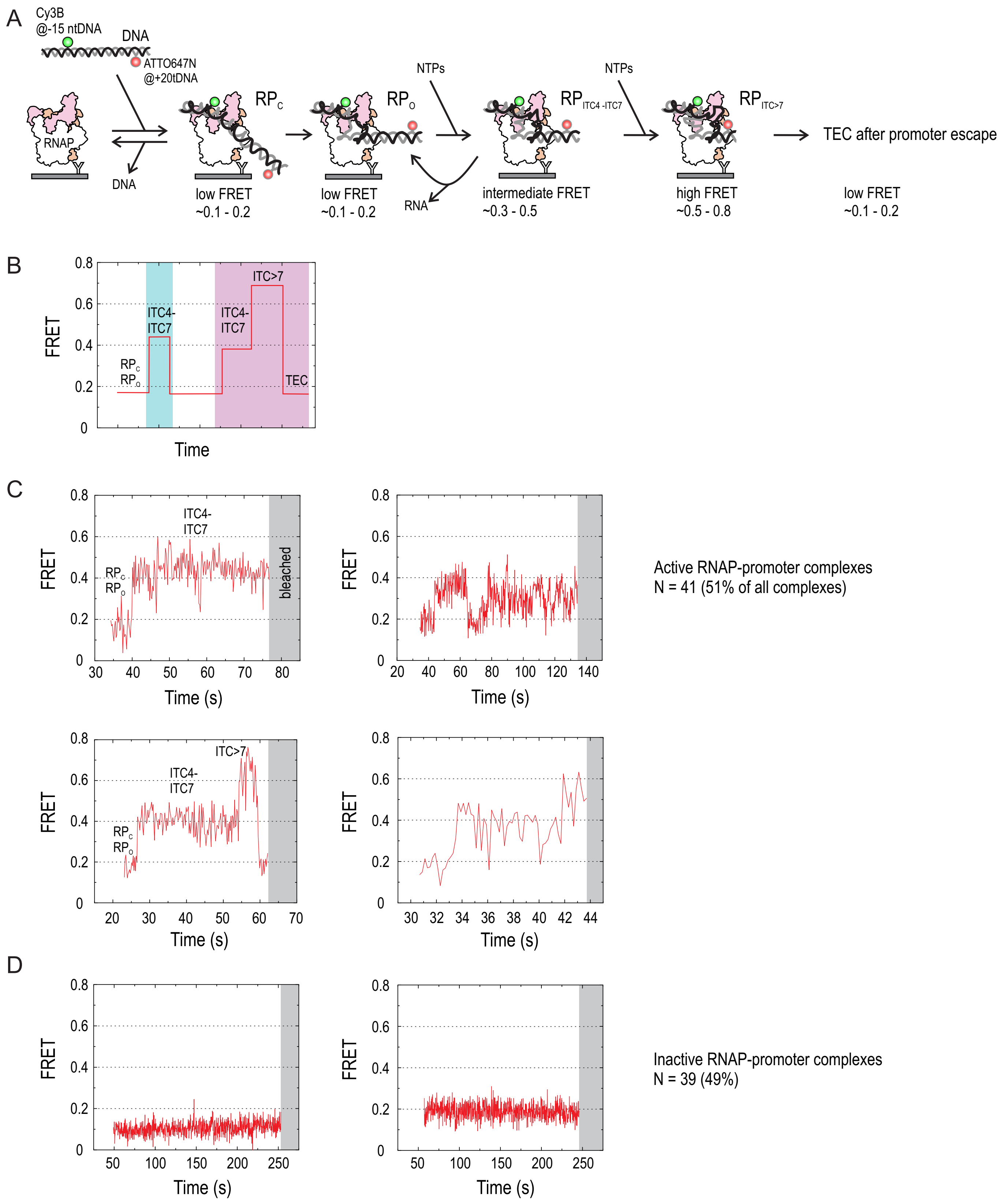
A
Cy3B
@-15 ntDNA
DNA
ATTO647N
@+20tDNA
RNAP
DNA
RPC
low FRET
~0.1 - 0.2
RPO
low FRET
~0.1 - 0.2
NTPs
RNA
RPITC4 -ITC7
intermediate FRET
~0.3 - 0.5
NTPs
RPITC>7
high FRET
~0.5 - 0.8
TEC after promoter escape
low FRET
~0.1 - 0.2
B
FRET
Time
RPC
RPO
ITC4-
ITC7
ITC>7
TEC
C
FRET
Time (s)
bleached
Active RNAP-promoter complexes
N = 41 (51% of all complexes)
D
Inactive RNAP-promoter complexes
N = 39 (49%)

### Figure S4

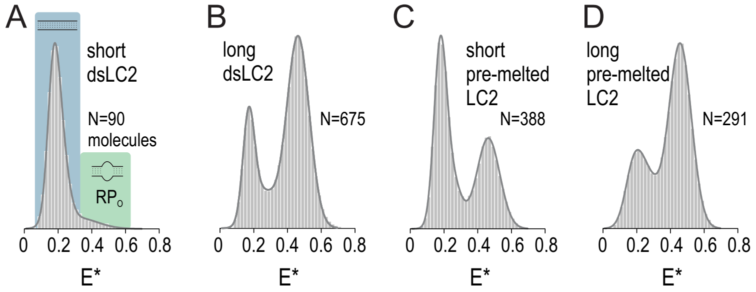
A
short
dsLC2
N=90
molecules
RPo
B
long
dsLC2
N=675
C
short
pre-melted
LC2
N=388
D
long
pre-melted
LC2
N=291
0 0.2 0.4 0.6 0.8
E*

Figure S5

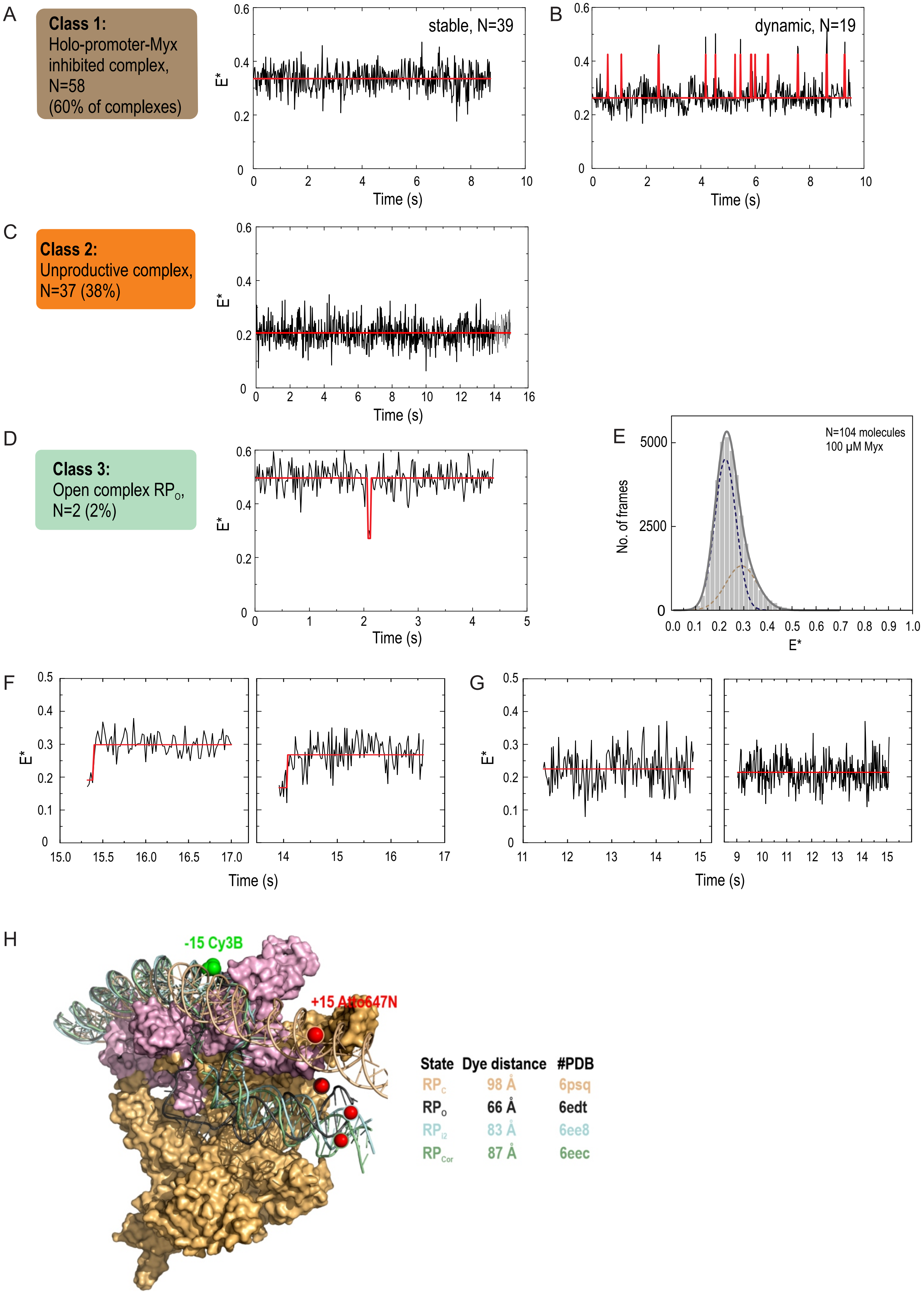
A
Class 1:
Holo-promoter-Myx inhibited complex, N=58
(60% of complexes)
stable, N=39
B
dynamic, N=19
C
Class 2:
Unproductive complex, N=37 (38%)
D
Class 3:
Open complex $RP_O$, N=2 (2%)
E
N=104 molecules
100 μM Myx
No. of frames
E*
F
G
Time (s)
H
-15 Cy3B
+15 Atto647N
State Dye distance #PDB
$RP_C$ 98 Å 6psq
$RP_O$ 66 Å 6edt
$RP_{I2}$ 83 Å 6ee8
$RP_{Cor}$ 87 Å 6eec

### Figure S6:

A

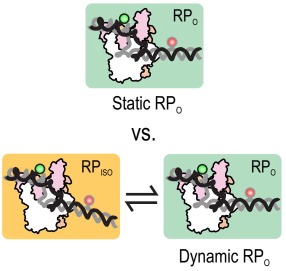


B

| | stable RP$_O$ | dynamic RP$_O$ |
|---|---|---|
| short dsLC2 | 76.8% (*N*=334 molecules) | 23.2% (101) |
| long dsLC2 | 85.3% (297) | 14.7% (51) |

molecules pooled from three independent experiments

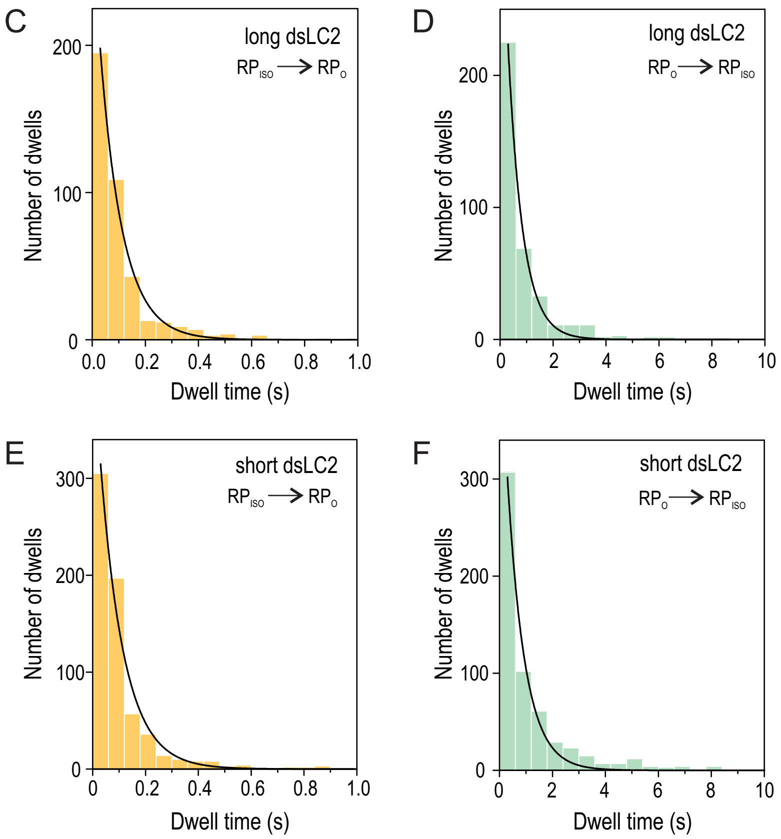


G

| | RP$_{ISO}$ → RP$_O$ | $\tau_{RPiso}$ (s) | RP$_O$ → RP$_{ISO}$ | $\tau_{RPo}$ (s) |
|---|---|---|---|---|
| short dsLC2 | | 0.090±0.006 | | 0.663±0.041 |
| long dsLC2 | | 0.085±0.004 | | 0.559±0.027 |
